# Neural Decoding and Feature Selection Techniques for Closed-Loop Control of Defensive Behavior

**DOI:** 10.1101/2024.06.06.597165

**Authors:** Jinhan Liu, Rebecca Younk, Lauren M Drahos, Sumedh S Nagrale, Shreya Yadav, Alik S Widge, Mahsa Shoaran

## Abstract

**Objective:** Many psychiatric disorders involve excessive avoidant or defensive behavior, such as avoidance in anxiety and trauma disorders or defensive rituals in obsessive-compulsive disorders. Developing algorithms to predict these behaviors from local field potentials (LFPs) could serve as foundational technology for closed-loop control of such disorders. A significant challenge is identifying the LFP features that encode these defensive behaviors.

**Approach:** We analyzed LFP signals from the infralimbic cortex and basolateral amygdala of rats undergoing tone-shock conditioning and extinction, standard for investigating defensive behaviors. We utilized a comprehensive set of neuro-markers across spectral, temporal, and connectivity domains, employing SHapley Additive exPlanations for feature importance evaluation within Light Gradient-Boosting Machine models. Our goal was to decode three commonly studied avoidance/defensive behaviors: freezing, bar-press suppression, and motion (accelerometry), examining the impact of different features on decoding performance.

**Main results:** Band power and band power ratio between channels emerged as optimal features across sessions. High-gamma (80-150 Hz) power, power ratios, and inter-regional correlations were more informative than other bands that are more classically linked to defensive behaviors. Focusing on highly informative features enhanced performance. Across 4 recording sessions with 16 subjects, we achieved an average coefficient of determination of 0.5357 and 0.3476, and Pearson correlation coefficients of 0.7579 and 0.6092 for accelerometry jerk and bar press rate, respectively. Utilizing only the most informative features revealed differential encoding between accelerometry and bar press rate, with the former primarily through local spectral power and the latter via inter-regional connectivity. Our methodology demonstrated remarkably low time complexity, requiring *<*110 ms for training and *<*1 ms for inference.

**Significance:** Our results demonstrate the feasibility of accurately decoding defensive behaviors with minimal latency, using LFP features from neural circuits strongly linked to these behaviors. This methodology holds promise for real-time decoding to identify physiological targets in closed-loop psychiatric neuromodulation.

## 1 Introduction

Fear and anxiety serve as adaptive defensive responses to threats, a phenomenon observed across a variety of species [1]. These responses are evolutionary mechanisms designed to enhance survival by preparing an organism to confront or flee from immediate danger [2, 3]. However, the same reactions, when excessive or misplaced, can significantly disrupt an individual’s overall quality of life [4, 5]. Anxiety disorders, characterized by disproportionate and persistent fear and anxiety, are among the most prevalent psychiatric conditions [6, 7]. Fear and anxiety contribute to the manifestation of a wide array of psychiatric disorders, underscoring their critical role in mental health [8, 9].

Fear and anxiety are often studied through the lens of defensive behaviors, a set of responses or patterns elicited in the face of perceived threats [10]. Rodents exhibit various defensive behaviors in response to actual or potential threats [11, 12]. These behaviors are often used to model human illnesses, because anxiety can also be viewed as an excessive response to potential or actual threats [11,13]. Therefore, exposure to a threatening stimulus evokes defensive responses that resemble emotional states related to fear and anxiety [12]. Recent studies indicate that the defensive patterns observed in normal human subjects show notable similarities to those of laboratory rodents. This parallel supports the hypothesis that rodent defensive behaviors may be reasonable models of similar behaviors in human anxiety disorders [14, 15]. The subjective human experience of fear is not the same as innate defensive behaviors in lower vertebrates, but those defensive behaviors are the closest available model [16]. Furthermore, both human fear/anxiety and rodent defensive behavior load onto the same frontal-amygdala circuits [17–19]. Hence, animal defensive behaviors offer a valuable model for understanding negative-valence processes in humans [20–22]. As a result, the excessive or contextually inappropriate exhibition of these behaviors can serve as a model for certain aspects of human psychiatric disorders [23, 24].

The long-term goal of modeling defensive and anxious behavior is to develop new treatments. Direct electrical stimulation of the brain is a particularly promising approach to that translation. Brain stimulation specifically improves the symptoms of multiple fear/anxiety disorders [25–28]. The approach involves the precise targeting of specific brain areas to modulate dysfunctional neural circuits associated with these conditions, which allows direct targeting of mechanisms discovered through animal models. Within the field of psychiatric brain stimulation, there is a strong drive towards closed-loop therapies [29–32]. The symptoms of psychiatric disorders, and of fear/anxiety disorders in particular, vary over time, and only some of those symptom states require neurostimulation. A closed-loop brain-machine interface (BMI) system that uses real-time neural activity from the subject to guide stimulation could help develop effective, precisely tailored therapies that stimulate only when it will be beneficial [29, 33–36].

There exists a main challenge for developing such closed-loop BMIs: we need a neural decoder that is capable of estimating the disorder symptom or behavior in real-time [37–39]. The development of highly accurate and fast decoders is essential for optimizing the therapeutic outcomes, ensuring that stimulation protocols are dynamically adjusted to the fluctuating patterns of neural dysregulation associated with psychiatric disorders [40]. Consequently, advancements in BMI technology and decoding algorithms hold the promise of revolutionizing the treatment landscape for patients with psychiatric conditions, offering hope for more personalized and effective interventions.

Decoding plays a pivotal role in neural engineering and the analysis of neural data [41–43]. It leverages activity recorded from the brain to forecast behaviors or symptoms [44–47]. These predictions, derived from decoding, can be utilized to manipulate devices or to enhance our understanding of the brain’s involvement in disorders [48–50]. This is achieved by assessing the extent of information that neural activity conveys about a symptom or behavior and examining how this information varies across different brain areas, experimental conditions, and states of disorder [51–53]. Decoding psychiatric states poses unique modeling challenges due to the complex and widespread network of brain regions involved in neural processes linked to neuropsychiatric states and behaviors, particularly in disorders such as chronic pain, addiction, or post-traumatic stress disorder (PTSD) [30, 38, 54–57].

In essence, neural decoding represents a regression or classification challenge that links neural signals to specific variables [58]. Machine Learning (ML) has emerged as a pivotal technique for elucidating the intricate patterns of neural activity, as well as the individual variations in brain function correlated with symptoms and behaviors [59]. Its utility is particularly pronounced when the primary research objective is to enhance predictive accuracy, a goal partly attributed to ML’s proven efficacy in addressing nonlinear challenges [60, 61]. Despite recent advances in ML techniques, the decoding of neural activity frequently employs traditional approaches such as linear regression (LR) and support vector machine (SVM) [62–64]. The adoption of contemporary ML tools for neural decoding might yield not only a substantial performance improvement but also the possibility of gaining more profound insights into neural functionality, as shown in recent studies. Encoding-decoding frameworks, based on linear state-space models, have decoded mood and cognitive state fluctuations from multisite intracranial electrocorticogram (ECoG) or stereo-electroencephalography (sEEG) signals [38, 62]. A Multi-layer Perceptron (MLP) has been utilized to forecast depressive states in human patients from local field potential (LFP) signals [65]. A decoder leveraging Random Forest models has been developed for the prediction of multi-class affective behaviors via intracranial electroencephalography (iEEG) recordings from the human mesolimbic network [66]. A discriminative cross-spectral factor analysis model was utilized for identifying a brain-wide oscillatory pattern for predicting resilient versus susceptible mice to stress [67]. Episodes of mental fatigue and changes in vigilance were precisely decoded from ECoG signals in Non-Human Primates (NHPs), using a gradient boosting classifier [68, 69].

In the aforementioned studies, beyond the decoding model utilized for prediction, the neuro-markers derived from neural data were crucial for decoding efficacy. The majority of previous efforts to detect psychiatric symptoms and behaviors have focused on classical spectral power features [38, 62, 66, 70, 71]. It is not clear that spectral power is the best feature for decoding complex cognitive-emotional phenomena such as fear/defensive behaviors. For instance, spectral power features were outperformed by cross-region connectivity metrics when attempting to decode cognitive task engagement [63, 72]. Spectral (wavelet entropy), temporal (Hjorth parameters), and connectivity features (partial directed coherence and phase locking index) have all been identified as significant markers for detecting mental fatigue [68]. In contrast, shifts in depressive states were more influenced by variations in spectral power features within the subcallosal cingulate than by coherence and phase-amplitude coupling [65]. Consequently, the importance of spectral power vs. other neuro-markers for modeling and decoding defensive behaviors and fear expression requires further investigation.

Here, we studied the decoding of defensive behaviors from the prefrontal cortex (PFC) and amygdala, which together comprise a circuit believed to regulate the expression of threat/defense versus safety behaviors [11, 24, 73, 74]. In prior rodent work, the balance between defensive and safety behaviors was associated with theta band (5-8 Hz) LFP synchrony between the infralimbic cortex (IL) and basolateral amygdala (BLA) [75–77]. Therefore, IL-BLA LFP connectivity and power features are promising targets for the development and testing of decoding algorithms that could be used in closed-loop psychiatric BMIs. At the same time, prior work focused on simple categorical analyses (t-tests between groups) and did not consider the more clinically relevant question of how to decode imminent behavior at the timescale of milliseconds to seconds. Rapid decoding would be crucial for a closed-loop BMI aimed at mitigating anxious or avoidance behavior in humans. It is not clear that the same LFP features that broadly discriminate two groups will be able to predict moment-to-moment behavior.

We thus developed a behavioral decoder based on IL-BLA LFP signals from rats undergoing a tone-shock conditioning and extinction protocol [78, 79]. Beyond conventional band power features, we explored and exploited a broad array of neuro-markers derived from the LFPs, across spectral, temporal, and connectivity domains. We considered three defensive behaviors: freezing, bar press suppression (bar press rate), and accelerometry, specifically the jerk (first derivative) calculated from a 3-axis head-mounted accelerometer. Freezing and bar press suppression are canonical defensive behaviors that have been studied for decades [20–22, 80–82]. Accelerometry jerk is a newer metric we have proposed and shown to correlate with, but not fully overlap the two other measures [83]. We developed a decoding framework based on Light Gradient-Boosting Machine (LightGBM), which outperformed other state-of-the-art ML-based decoders in both accuracy and latency. Our approach included a methodology to assess feature importance and a feature selection strategy utilizing the SHapley Additive exPlanations (SHAP), effectively reducing the dimensionality of the feature space. Band power and band power ratio between channels emerged as critical for decoding defensive behaviors, with the high-gamma band being particularly predictive compared to other frequency bands. By prioritizing highly-ranked neuro-markers, we enhanced decoding performance beyond that with solely band power features. Consequently, this study underscores the effectiveness of our proposed ML framework in the precise and rapid decoding of defensive behaviors within a closed-loop psychiatric Brain-Machine Interface (BMI) system.

## 2 Methods

### 2.1 Animals and behavior paradigm

We utilized 16 adult Long Evans rats, with weights ranging from 250 to 350 grams. Initially, rats were pair-housed in plastic cages for at least 7 days to facilitate acclimation to the research facility. Subsequent to this acclimation period, the rats underwent daily handling for 5 days to mitigate handling-related stress, after which they were individually housed in plastic cages. To prepare for experimental procedures, food intake was restricted to 10 grams per day until each rat achieved 85-95% of its initial body weight. Thereafter, the animals were allocated 15-20 grams of food daily to maintain their weight within this specified range throughout the behavioral experiments. During the first three days of food restriction, sucrose pellets were introduced into the home cages to acquaint the rats with the reward, thereby facilitating the learning of bar-pressing behavior.

The behavioral training and experiments were conducted in the Coulbourn conditioning chambers, with dimensions of 30.5 *×* 24.1 *×* 21 cm. These chambers were equipped with a grid floor, consisting of rods spaced 1.6 cm apart and with a diameter of 4.8 mm, to facilitate the delivery of foot shocks. An aluminum wall of the chamber incorporated a retractable bar and a food trough for monitoring reward-seeking behaviors, while a speaker mounted on the opposite wall emitted sound stimuli. Additionally, a camera with an attached wide-angle lens was positioned outside the conditioning chamber, above the speaker unit, to record video footage through the chamber’s plexiglass top.

Initially, rats were trained to execute bar presses to obtain sucrose pellets. They subsequently underwent electrode implantation and participated in a post-surgical behavioral paradigm. To provoke defensive behaviors, the rats were exposed to a tone-shock conditioning protocol, which comprised three phases: habituation/conditioning, extinction, and extinction recall, as shown in Figure 1(a). Electrophysiological and video recordings were systematically carried out during each experimental session. During the habituation phase on day 1, rats encountered 5 trials of the conditioned stimulus (CS: a 30-second, 82 dB tone). This was followed by the conditioning phase, where they experienced 7 instances of the CS paired with the unconditioned stimulus (US: a 0.6 mA, 0.5-second foot shock) immediately after the CS. On day 2, the extinction phase consisted of 20 presentations of the CS alone, without the US, in the same chamber. On day 3, to evaluate extinction memory (recall), the CS was presented 6 times without the US.

**Figure 1.**
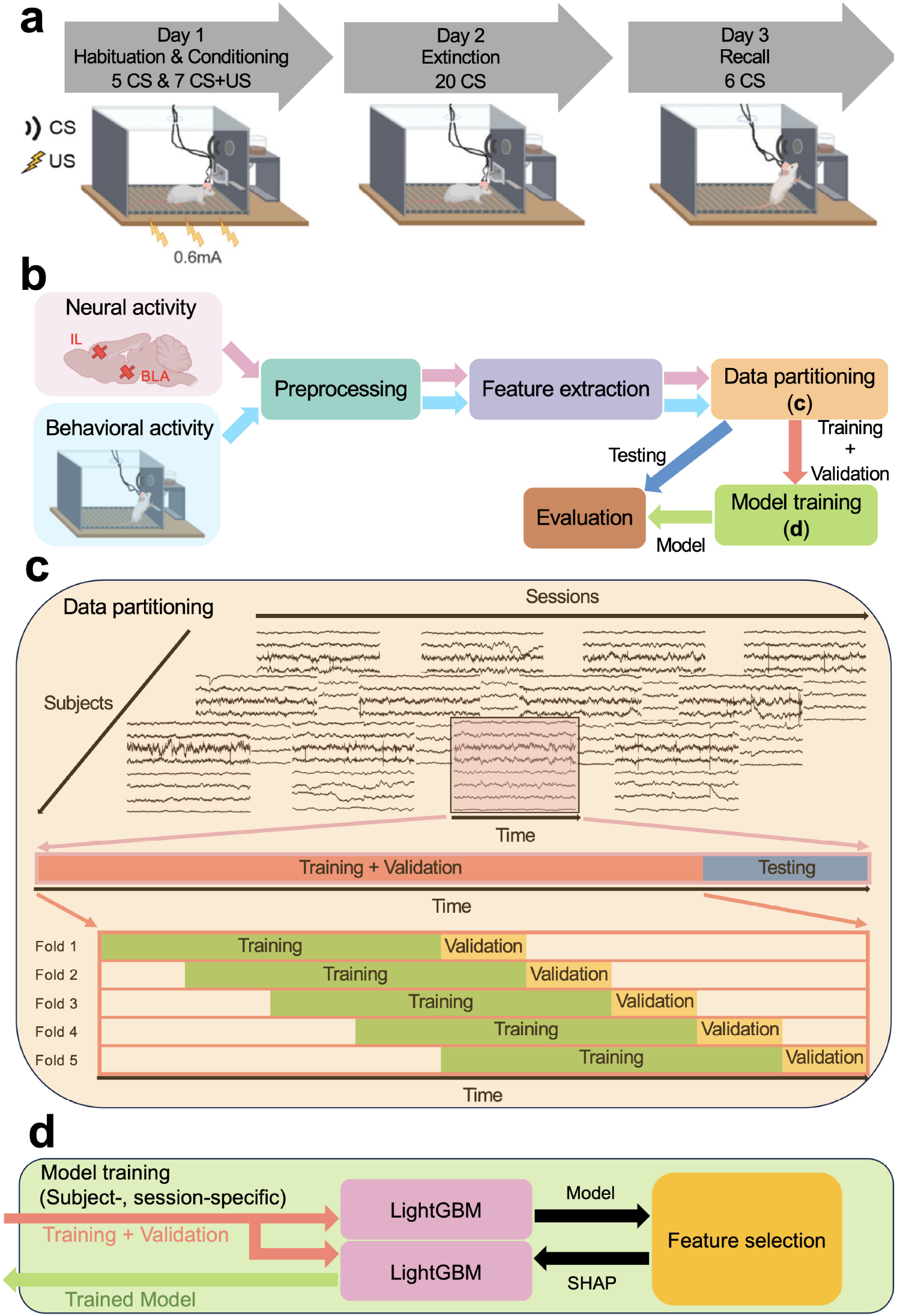
Experimental paradigm and proposed ML framework for decoding defensive behaviors. (a) The three-day tone-shock conditioning protocol. The experiment consisted of three phases: habituation/conditioning, extinction, and extinction recall, with electrophysiologic and behavioral data recorded during each phase. (CS: Conditioned stimulus. US: Unconditioned stimulus) (b) The proposed ML framework contained modules including neural data preprocessing, feature extraction in three representation domains, data partitioning into training, validation and testing sets as shown in (c), decoding model training as shown in (d), and model evaluation. (c) Data partitioning process in each recording session for each subject. A hold-out testing set was taken at the end of the session, and the beginning of the session was split in a sliding window 5-fold crossvalidation paradigm. (d) The procedure of training the decoding model. SHAP values were measured using the first trained LightGBM model with all features, and the second LightGBM was then trained using the high-rank features in the order of their SHAP values and subsequently used for final model evaluation.

Reward-seeking behavior, indicated by bar presses, functioned as a dynamic measure for defensive behavior, with a decrease in pressing activity interpreted as an elevated threat response. Bar press events were captured in the electrophysiology event data using a Data Acquisition System (DAQ) (USB 6343-BNC or PCIe-6353, National Instruments, Woburn, MA, USA). These event data were subsequently processed to isolate bar press incidents and their associated timestamps. Assessment of freezing behavior was conducted through offline video analysis, employing a Logitech HD Pro Webcam C910 equipped with a Neewer Digital High Definition 0.45x Super Wideangle Lens. The footage, captured at a rate of 24 frames per second with Debut Professional software, was analyzed using ANY-maze, which utilizes its integrated freezing detection functionality to assign a ‘freezing score’ to each frame. This score increased with more significant changes in pixels between consecutive frames. Meanwhile, accelerometry data were collected continuously at a 30 kHz sampling rate via the RHD 2132 electrophysiology headstage, which includes a built-in 3-axis accelerometer. These data were logged using the Open Ephys acquisition system, a widely used open-source platform for in-vivo electrophysiology research [84]. The synchronization of accelerometry records with video data was accomplished by aligning the ‘tone on’ events observed in both datasets. There were sparse bar press events for a few rats during the conditioning phase due to the bar press suppression resulting from foot shocks. We chose 10 rats with no less than 5 bar presses in the conditioning phase to ensure the threat responses were well-encoded in the following neural decoding study. All 16 rats were used for the analysis in habituation, extinction, and recall sessions.

### 2.2 Electrodes and surgery

Each electrode bundle was comprised of 8 nickel-chromium recording microwires, each measuring 12.5 *µ*m in diameter, accompanied by one reference wire of the same diameter as the recording wires, and a platinum-iridium stimulating channel with a diameter of 127 *µ*m [85]. The stimulation channel was used for another set of experiments not reported here, and no stimulation occurred during any of the data reported in this paper. These recording and stimulating components were collectively bundled within a 27-gauge stainless steel cannula, which also served as a pathway for the return current during stimulation. The recording wires were bonded to the individual pins of an Omnetics connector using silver paint, while the stimulating wire was soldered to a mill-max connector, enabling concurrent recording and stimulation at the same site. A ground wire was affixed to the connector, and the entire bundle was safeguarded with epoxy. Prior to surgery, the electrodes were sterilized using Ethylene Oxide (EtO).

The electrode arrays were surgically implanted into the left infralimbic cortex (IL) (+3 mm anterior-posterior (AP), +0.5 mm medial-lateral, and -3.95 mm dorsal-ventral (DV) from the brain surface) and the basolateral amygdala (BLA) (-2.28 mm AP, +5 mm medial-lateral, and -7.5 mm DV from the brain surface). Dental cement was utilized to secure the electrodes and to construct a protective head cap for the animals. The ground wire was securely wrapped around a skull screw prior to the implant fixation. A minimum recovery period of seven days was allowed for the animals before starting physiological experiments.

### 2.3 Electrophysiology and data processing

The electrophysiological signals, specifically local field potentials (LFPs), were recorded at a sampling rate of 30 kHz using an Open Ephys acquisition system throughout all experimental sessions. The recording headstage was interfaced with two mill-max male-male connectors, each comprising eight channels, through an adaptor.

Quantification of defensive behavior adhered to the methodology established in a prior study [83]. The change in total acceleration, herein referred to as “accelerometry jerk”, was computed as:

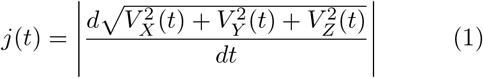

where *j*(*t*) is the accelerometry jerk as a function of time *t*, and 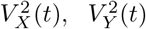, and 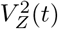 are the voltages of the accelerometer in *X, Y* , and *Z* axes, respectively. The accelerometry jerk was downsampled from its original sampling rate of 30k samples/second to 1k samples/second, and was then smoothed with a Gaussian filter using a 200-sample window to remove non-biological noise transients. Bar press events and their corresponding timestamps were extracted from the electrophysiological recordings, with timestamps being resampled to 1k samples/second. The counts of presses was binned into each 1-ms time interval, and then these counts were smoothed using a Gaussian filter with a 1k-sample window. This process transformed the data from a discrete count of events into an approximation of a continuous press rate, hereby referred to as the bar press rate. The resampling process utilized the *downsample*() function in Matlab, while Gaussian smoothing was executed with the *smoothdata*() function in Matlab.

A total of 8 recording channels were obtained from bipolar-referenced LFP signals, with 4 channels from each of the IL and BLA regions. These bipolar-subtracted channels were subsequently band-pass filtered within the range of 1-150 Hz using a 3rd-order zero-phase Infinite Impulse Response (IIR) Butterworth filter. Subsequently, line noise was removed by applying a notch filter at 60 Hz and its harmonics. The LFP data was then demeaned across the time series for each channel. For each subject and recording session, we visually inspected the neural data and excluded time epochs that exhibited clear non-neural artifacts, such as significant sharp voltage transients. For extracting features in various frequency bands, the neural data were processed using 3rd-order zero-phase IIR Butterworth band-pass filters across 7 frequency bands: 1-4 Hz (delta), 4-8 Hz (theta), 8-13 Hz (alpha), 13-30 Hz (beta), 30-50 Hz (low-gamma), 50-80 Hz (gamma), and 80-150 Hz (high-gamma). Phase and amplitude were extracted from the band-pass filtered signal via Hilbert transform. Cross-spectral density was estimated on the neural signals before band-pass filtering using the Multitaper method in the MNE package. The other preprocessing steps were implemented using the SciPy package.

Both behavioral data and the neuro-markers were computed in overlapping 1-second sliding windows with a 0.2-second step size. Behavioral measurements were quantified by averaging the measures of accelerometry jerk, bar press rate, and freezing score within each window.

### 2.4 Neuro-marker extraction

To investigate the neural representations in various aspects and enhance the accuracy of decoding defensive behaviors in our model, we extracted 17 types of neuro-markers across 3 representation domains as neural features for each window, as detailed in Table 1. We chose these features based on existing evidence that, in general, local power and cross-region connectivity between IL and BLA have been linked to defensive behaviors in past research [75, 86]. Additionally, we computed time domain features that are pivotal in identifying patterns of neural activity associated with specific behaviors or pathological states, owing to their simplicity and the direct interpretation of neural dynamics [87–89].

**Table 1:**
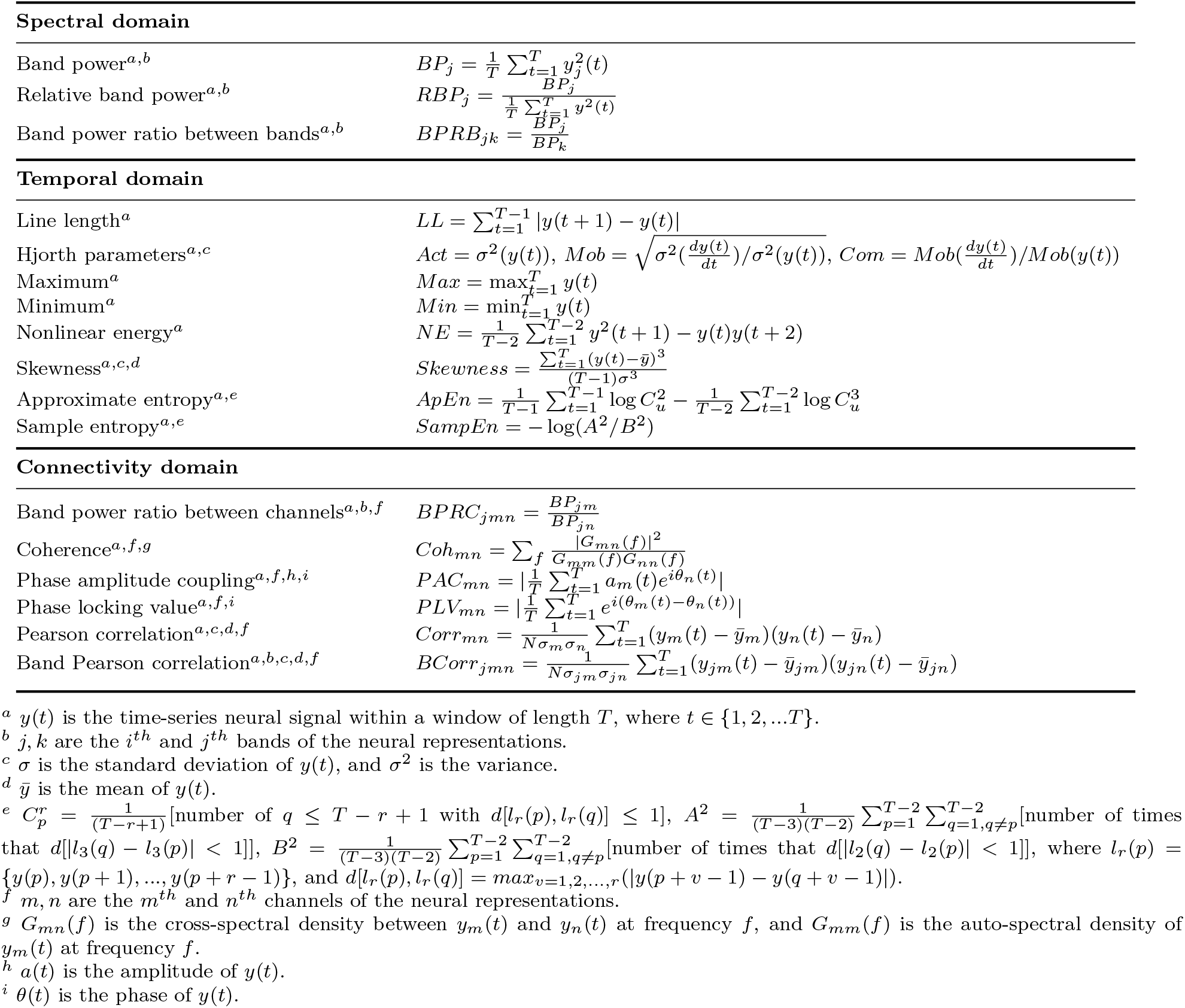
Neuro-markers extracted in the spectral, temporal, and connectivity domains.

In the spectral domain, band power (BP) was quantified across 7 frequency bands [38, 90, 91]. Relative band power (RBP) refers to the power in a specific frequency band relative to the total signal power [87, 92, 93]. Band power ratio between bands (BPRB) facilitates the pairwise comparison of power levels across different bands within a single channel [92–94].

Regarding temporal features, line length (LL) calculates the absolute differences between successive time points [87, 90, 95]. Hjorth parameters (HP) reflect statistical attributes including variance, mean frequency, and frequency variation [87, 96–98]. Maximum (Max) and minimum (Min) represent the extreme values within the window [87]. Nonlinear energy (NE) gives an estimate of the energy content of the neural signal [99]. Skewness evaluates the asymmetry of the distribution of instances within the window [100]. Approximate entropy (ApEn) and sample entropy (SampEn) assess the existence of patterns within a sequence of instances [101, 102].

In the connectivity domain and cross-region representations, band power ratio between channels (BPRC) enables pairwise power level comparison across channels from two distinct brain regions [92]. Coherence (Coh) quantifies the similarities of neural oscillation between channels [103]. Phase-amplitude coupling (PAC) captures the linkage between the phase of a low-frequency band and the amplitude of a high-frequency band between channels [88, 104, 105]. Phase locking value (PLV) describes the phase relationship consistency between signals from different channels [77, 106]. Pearson correlation (Corr) and band Pearson correlation (BCorr) quantify functional connectivity between channels, across the full band and within individual frequency bands, respectively [72].

Here, BP, RBP, BPRC, Coh, PLV, and BCorr were assessed across the aforementioned 7 frequency bands. PAC analysis was performed between the amplitudes of the low-gamma, gamma, and high-gamma bands and the phases of the theta and alpha bands, with 6 phase-amplitude combinations. BPRB comparisons were made between each pair of the 7 bands, with 21 band-band combinations in total. BP, RBP, BPRB, LL, HP, Max, Min, NE, skewness, ApEn, and SampEn were calculated for each individual channel. BPRC, Coh, PAC, PLV, Corr, and BCorr were derived only from channel pairs between IL and BLA, with 16 channel-channel combinations. ApEn and SampEn were computed using the MNE-Features package.

### 2.5 Data partitioning

After extracting the neuro-markers and behavioral data, we partitioned them into three distinct datasets for subsequent use in training the decoding model, selecting high-rank features, and evaluating the model’s performance. For each subject, we divided the data from each recording session into training, validation, and test sets, as depicted in Figure 1(c).

A hold-out test set, constituting 20% of the entire recording, was designated from the final 20% of each behavior session, while the initial 80% served as the training and validation sets. The separation between the training and validation sets employed a sliding-window 5-fold cross-validation paradigm. The time series data were evenly divided into 9 windows. In the first fold, the initial 4 windows formed the training set, and the 5th window served as the validation set. From the second to fifth folds, we sequentially shifted the training and validation sets by one window forward in time, ensuring that validation sets were different across folds and incorporating validation sets from preceding folds into the training sets of subsequent folds. Therefore, the division ratios for training and validation sets versus test sets, and training sets versus validation sets, were maintained at 80%-20%. This method respected the chronological sequence of the time series data by consistently organizing the datasets in a training-validation-testing order. This organization assured that the model was always trained on historical data and validated/tested on subsequent data, thereby preventing data leakage across the temporal dimension.

The test set was utilized for the final evaluation of the model’s decoding accuracy, trained using the complete training and validation sets. Feature selection and parameter optimization were conducted based on the model’s validation set performance, trained on the training set data.

### 2.6 Decoding model

A diverse array of machine learning (ML) models has been employed for neuropsychiatric tasks and brain-machine interface applications, including linear regression (LR) [62, 103], support vector machine (SVM) [63, 67, 77, 107, 108], random forest (RF) [109, 110], and artificial neural network (ANN) [111, 112]. Moreover, gradient-boosted decision trees (GBDT) have demonstrated promising performance in previous neurophysiological task studies [90, 91, 97, 107, 113]. In this work, we utilized a GBDT-based model named Light Gradient-Boosting Machine (LightGBM), known for its efficiency in reducing data instances and features through gradient-based one-side sampling (GOSS) and exclusive feature bundling (EFB) [114]. GOSS retains data instances with gradients above a certain threshold while randomly discarding instances with smaller gradients, thereby maintaining the data’s substantial contribution to information gain. EFB efficiently reduces the number of effective features by bundling mutually exclusive features — those not taking non-zero values simultaneously — into a single feature. By leveraging GOSS and EFB, LightGBM achieves superior computational speed and lower memory usage compared to other GBDTs, without compromising the accuracy intrinsic to GBDT models. We configured LightGBM with 100 trees, setting the maximum number of leaves per tree to 5 to prevent overfitting. These parameters were determined through hyperparameter optimization during the cross-validation phase.

In addition to LightGBM, we evaluated a variety of ML models widely applied in neurophysiological research, employing our proposed feature set as outlined in Table 1. These models encompass traditional ML approaches such as LR, SVM with both linear (SVM-Lin) and radial basis function kernels (SVM-RBF), the tree-based RF model, and ANN models with diverse architectures, including a 3-layer multilayer perceptron (MLP), 3-layer long short-term memory (LSTM), and 3-layer convolutional neural network (CNN). The implementation of LR, SVM-Lin, SVM-RBF, and RF was conducted using the scikit-learn package, while MLP, LSTM, and CNN were implemented via the Pytorch package. The LightGBM model was implemented using the Python package provided by Microsoft. In our preliminary decoding analysis, leveraging all neural features depicted in Table 2, we assessed the decoding accuracy for accelerometry jerk and bar press rate across the aforementioned ML models, averaged over subjects in each recording session. Performance evaluation was conducted using both the coefficient of determination (*R*^2^) and the Pearson correlation coefficient (*r*) metrics. It should be noted that here, *R*^2^ is not the squared Pearson correlation coefficient, and its value lies within the range of (*−*∞, 1]. A negative *R*^2^ suggests that the decoded behavior captures less variation in the real behavior than a constant value equivalent to the average of the ground truth, indicating relatively poor decoding performance. Our findings indicate that LightGBM outperformed other ML models in decoding both accelerometry jerk and bar press rate in 14 out of 16 comparisons.

**Table 2:**
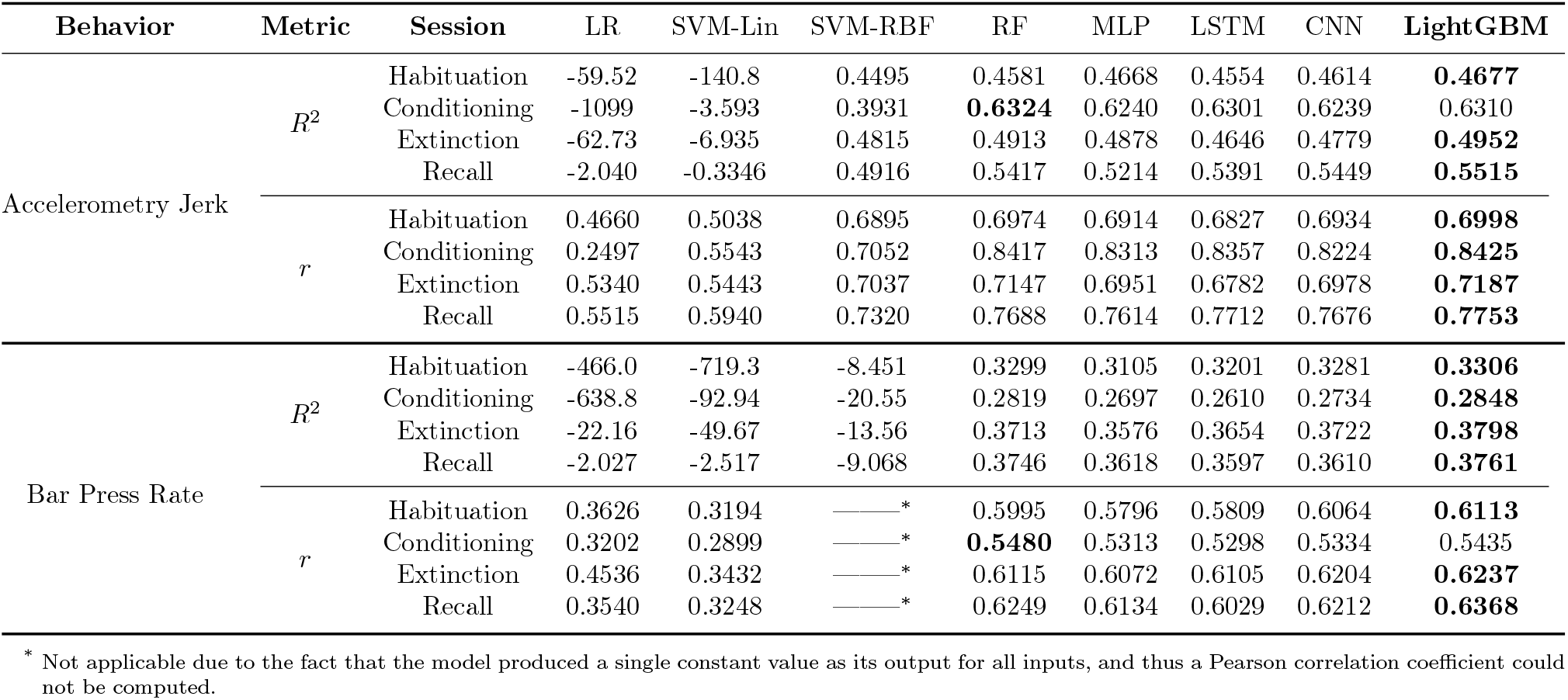
Performance of ML models for decoding defensive behaviors averaged across subjects in each recording session. The performance was evaluated using the coefficient of determination (*R*^2^) and the Pearson correlation coefficient (*r*). The best results are **bolded**.

| Behavior | Metric | Session | LR | SVM-Lin | SVM-RBF | RF | MLP | LSTM | CNN | LightGBM |
| --- | --- | --- | --- | --- | --- | --- | --- | --- | --- | --- |
| Accelerometry Jerk | $R^2$ | Habituation | -59.52 | -140.8 | 0.4495 | 0.4581 | 0.4668 | 0.4554 | 0.4614 | **0.4677** |
|  |  | Conditioning | -1099 | -3.593 | 0.3931 | **0.6324** | 0.6240 | 0.6301 | 0.6239 | 0.6310 |
|  |  | Extinction | -62.73 | -6.935 | 0.4815 | 0.4913 | 0.4878 | 0.4646 | 0.4779 | **0.4952** |
|  |  | Recall | -2.040 | -0.3346 | 0.4916 | 0.5417 | 0.5214 | 0.5391 | 0.5449 | **0.5515** |
| | $r$ | Habituation | 0.4660 | 0.5038 | 0.6895 | 0.6974 | 0.6914 | 0.6827 | 0.6934 | **0.6998** |
|  |  | Conditioning | 0.2497 | 0.5543 | 0.7052 | 0.8417 | 0.8313 | 0.8357 | 0.8224 | **0.8425** |
|  |  | Extinction | 0.5340 | 0.5443 | 0.7037 | 0.7147 | 0.6951 | 0.6782 | 0.6978 | **0.7187** |
|  |  | Recall | 0.5515 | 0.5940 | 0.7320 | 0.7688 | 0.7614 | 0.7712 | 0.7676 | **0.7753** |
| Bar Press Rate | $R^2$ | Habituation | -466.0 | -719.3 | -8.451 | 0.3299 | 0.3105 | 0.3201 | 0.3281 | **0.3306** |
|  |  | Conditioning | -638.8 | -92.94 | -20.55 | 0.2819 | 0.2697 | 0.2610 | 0.2734 | **0.2848** |
|  |  | Extinction | -22.16 | -49.67 | -13.56 | 0.3713 | 0.3576 | 0.3654 | 0.3722 | **0.3798** |
|  |  | Recall | -2.027 | -2.517 | -9.068 | 0.3746 | 0.3618 | 0.3597 | 0.3610 | **0.3761** |
| | $r$ | Habituation | 0.3626 | 0.3194 | —* | 0.5995 | 0.5796 | 0.5809 | 0.6064 | **0.6113** |
|  |  | Conditioning | 0.3202 | 0.2899 | —* | **0.5480** | 0.5313 | 0.5298 | 0.5334 | 0.5435 |
|  |  | Extinction | 0.4536 | 0.3432 | —* | 0.6115 | 0.6072 | 0.6105 | 0.6204 | **0.6237** |
|  |  | Recall | 0.3540 | 0.3248 | —* | 0.6249 | 0.6134 | 0.6029 | 0.6212 | **0.6368** |

\* Not applicable due to the fact that the model produced a single constant value as its output for all inputs, and thus a Pearson correlation coefficient could not be computed.

### 2.7 Model training and evaluation

Figure 1(d) illustrates the model training process using the dataset configuration detailed in Figure 1(c). LightGBM models were trained in a subject-specific and session-specific manner, premised on the hypothesis that neural representations of defensive behaviors exhibit inter-subject variability. Furthermore, we fitted models separately for each recording session because we expected the neural encoding to shift over time. Tone-shock conditioning and extinction learning both involve significant plasticity in the IL-BLA circuit, and thus defensive behaviors might be driven by different activity patterns before vs. after a given stage of learning. For each subject and session, 5 Light-GBM models were trained and assessed using the 5 folds designated for the training and validation sets, which were subsequently used for the selection of top-ranked features based on high feature importance values. A final LightGBM model was then trained using the aggregated training and validation sets and evaluated against the hold-out test set, incorporating either band power features, selected top-ranked features, or the entire set of extracted features.

The model’s decoding performance was quantified using *R*^2^ and *r* to compare ground truth with predicted behavioral measurements. The loss in *R*^2^ served to evaluate the neuro-markers’ contribution to decoding performance by their exclusion from the model, and it was also applied in the validation set’s performance analysis to guide the selection of a specific number of top-ranked features. Additionally, *r* was also utilized to compare the similarities between feature importance matrices. The evolution of the training curves, delineated by the percentage change in L2 Loss with increasing iterations, provided further insight into training dynamics.

### 2.8 Feature selection

Integrating an increased number of neuro-markers across spectral, temporal, and connectivity domains may enhance the decoding accuracy for defensive behaviors. However, this augmentation results in a proliferation of features, increasing computational complexity and memory requirements. Furthermore, some features may be uninformative or redundant within the ML framework, complicating the derivation of neuroscientific insights from models based on an extensive array of features. In our study, we extracted 17 types of neuro-markers, totaling 1296 features as input into the model, which inflated the computational costs unnecessarily. Consequently, we implemented a feature selection method to reduce computational demands, mitigate the risk of model overfitting, and identify which LFP features were most informative and, thereby, potentially causal to behavior.

We utilized SHapley Additive exPlanations (SHAP) for the assessment of feature importance among neuro-markers [115]. SHAP is a comprehensive measure of feature importance based on the Shapley values from a conditional expectation function of the original model. These values offer a unique feature importance metric that adheres to three desirable properties including local accuracy, missingness, and consistency when evaluating the additive attribution of one feature to the prediction output [115].

For each defensive behavior across every recording session, we assessed the SHAP values for every feature across the 5 LightGBM models, each trained using a distinct fold. Subsequently, for each feature, we computed the mean of its absolute SHAP values across all data instances and folds, establishing this as the cumulative contribution of the feature within that session. To identify the top-ranked features that are both subject- and behavior-specific and exhibit consistency across different days, these calculated attributions were further averaged over 4 recording sessions to determine the ultimate feature importance, as shown below:

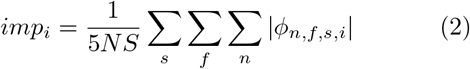

where *imp*_*i*_ is the importance of feature *i* ∈ {1, 2, …, *M*}, *ϕ*_*n*,*f*,*s*,*i*_ is the SHAP value for data instance *n* ∈ {1, 2, …, *N*}, fold *f* ∈ {1, 2, …, 5}, session *s* ∈ {1, 2, …, *S*}, and feature *i*, and *M, N, S* are the numbers of features, samples, and sessions, respectively. Then we sorted the feature importance *imp*_*i*_ and ranked the neural features for subsequent selection. We employed an iterative feature selection strategy, which means features were incrementally introduced to the model in the order of SHAP rankings. This process continued until the model’s performance on the validation set was comparable to, and not significantly lower than, the peak performance determined through an exhaustive iteration over an increasing amount of top-ranked features from one to the maximum. The group of features, when this process stopped, was the final selected subset of features used for decoding corresponding defensive behavior across recording sessions.

### 2.9 Statistical analysis

We conducted paired-sample t-tests to assess the differences in decoding performance between accelerometry jerk, bar press rate, and freezing score. Additionally, these tests compared the SHAP values of features without or with various temporal delays, by employing neural features not only from the current window, but also from the preceding windows lagged by up to 20 seconds, across all recording sessions and subjects. Because there is an inherent motor delay between perception of threat and emission of a defensive behavior, decoding might perform better if that delay were taken into account using this lagged method. We applied the same method to determine whether decoding performance using selected top-ranked features was significantly different from the optimal performance identified during feature selection.

Independent-sample t-tests were utilized to determine the statistical significance of overall SHAP feature importance within specific frequency bands relative to all other bands, across all recording sessions and subjects. This test was also applied to evaluate the significance of feature contributions to decoding performance within specific frequency bands in comparison with contributions from all other frequency bands. The Wilcoxon signed-rank test was employed to compare decoding performance when using band power features, selected top-ranked features, and the entire set of extracted features across all subjects. To account for multiple comparisons, Bonferroni corrections were applied to adjust p-values, tailored to the number of comparisons conducted. The implementation of paired-sample t-tests, independent-sample t-tests, and the Wilcoxon signed-rank tests were carried out using the SciPy package.

## 3 Results

### 3.1 Comparison of decodability of defensive behaviors using proposed ML framework

The comparison of training processes and decoding performances for accelerometry jerk, bar press rate, and freezing score is depicted in Figure 2. Figure 2(a) depicts the training curves, showcasing the L2 loss changes, averaged across subjects and recording sessions. The models underwent training using the training set, with the percentage change in L2 loss from the initial untrained state evaluated on both the training and validation sets. For all three behaviors, the L2 loss for training sets exhibited a consistent decline with additional iterations. However, the validation set loss for the freezing score demonstrated minimal improvement (-9.5%), in contrast to accelerometry jerk (-53.6%) and bar press rate (-34.6%).

**Figure 2.**
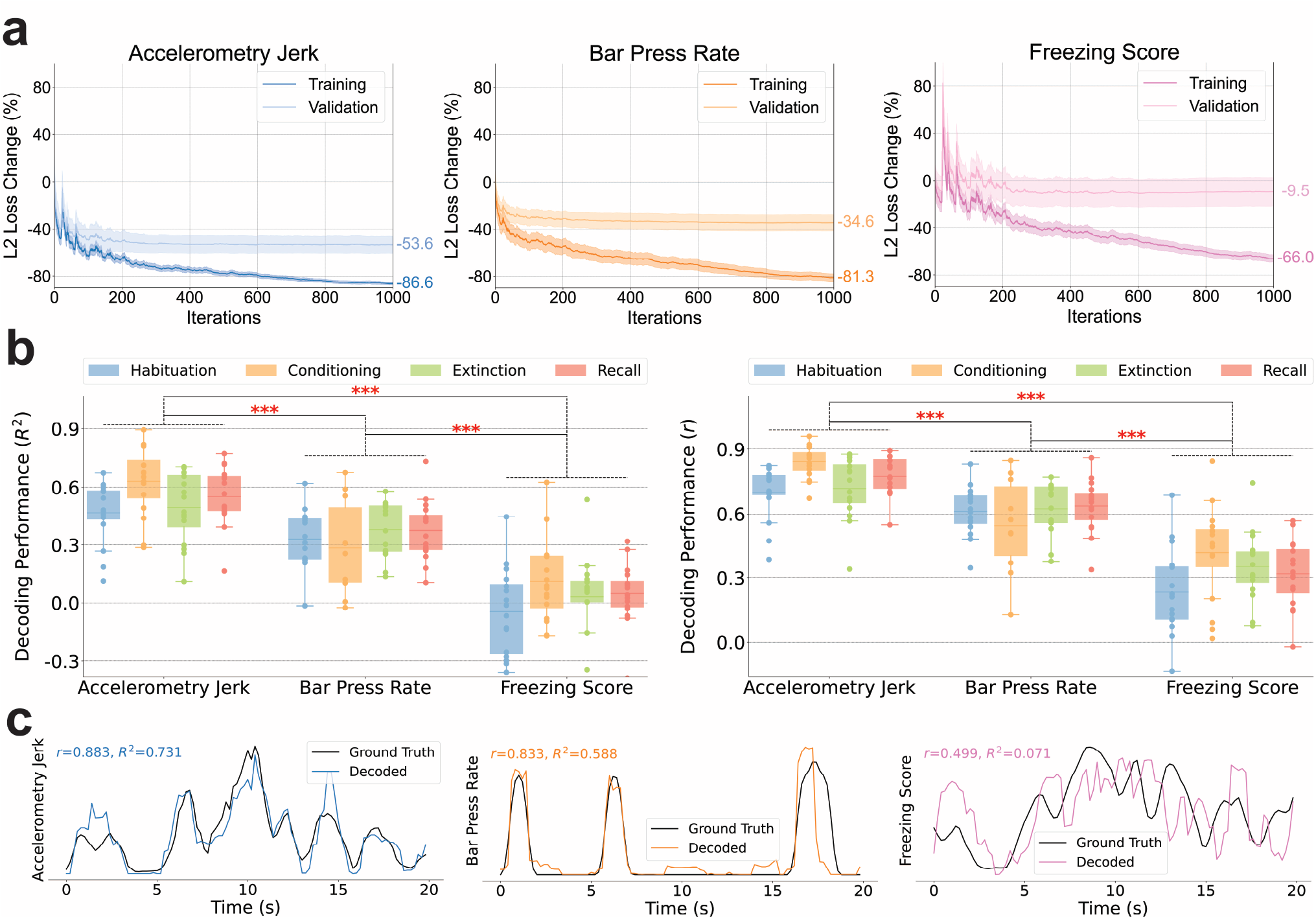
Decoding performances for accelerometry jerk, bar press rate, and freezing score. (a) Comparisons of training curves of our ML model for decoding accelerometry jerk (Left), bar press rate (Middle), and freezing score (Right). The percentages of the change of L2 Loss on the training and validation sets are shown with the increasing number of iterations during the training process. The shading areas indicate standard errors across subjects. (b) Decoding performance for accelerometry jerk, bar press rate, and freezing score in four recording sessions averaged across subjects. The line in the boxplot shows the average performance across subjects, and each dot indicates the result of an individual subject. The performances were evaluated using the coefficient of determination (Left, *R*^2^) and the Pearson correlation coefficient (Right, *r*). The asterisks denote the significant difference in the decoding performance of two defensive behaviors. (Paired-sample t-test; *\* * \** : *p* < 0.001) (c) Decoding examples for accelerometry jerk (Left), bar press rate (Middle), and freezing score (Right), from a single rat and session (OB44, Habituation), with performance evaluated using *R*^2^ and *r*.

There were large differences in the degree to which the different forms of defensive behavior could be decoded from the IL/BLA LFPs (i.e., in the degree to which these behaviors were encoded within the LFPs in that brain circuit). Specifically, the freezing score was only marginally decodable across sessions, with the mean coefficient of determination (*R*^2^) averaged across subjects never surpassing 0.12 in all recording sessions, as shown in Figure 2(b). While the bar press rate showed a higher degree of decodability, performances were slightly diminished during the conditioning session, attributed to strong bar press suppression resulting from foot shocks. Accelerometry jerk emerged as the most reliably decodable behavior, with the mean *R*^2^ values across subjects consistently exceeding 0.46 in all recording sessions. Overall, decoding accuracy varied significantly among different defensive behaviors, following a descending order from accelerometry jerk to bar press rate to freezing score. These findings remained consistent when evaluated using both *R*^2^ and the Pearson correlation coefficient (*r*) for performance assessment.

The variation in decoding performance may arise in part from the distinct characteristics of each behavioral signal, as illustrated in Figure 2(c). Accelerometry jerk is characterized by a smoothly fluctuating signal that remains predominantly nonzero. In contrast, bar press rate often drops to zero but then has sharp deviations from baseline during bouts of pressing. Freezing score exhibits some local deviations even after smoothing. Regarding the freezing score in Figure 2(c), the model succeeds in tracking the global trend, resulting in a relatively high *r*. Nevertheless, it struggles to capture local variations, leading to an *R*^2^ of 0.071 for the freezing score. This indicates that the decoded behavior scarcely captures variance from the actual behavior, offering only marginal predictive improvement over the expected value of the true behavior. In light of these findings, subsequent analyses concentrated on accelerometry jerk and bar press rate, given that interpretations derived from the non-predictive models of freezing score could potentially be misleading.

### 3.2 Importance and contribution of neuro-markers to the decoding performance

Subsequently, our focus shifted towards understanding the importance of each neuro-marker type in decoding defensive behaviors. Figures 3(a) and (b) delineate the importance of three feature domains, diverse neuro-markers, and frequency bands in decoding defensive behaviors. A notable observation is that spectral, temporal, and connectivity features all play a crucial role in decoding defensive behaviors. Specifically, temporal (43.0%) and spectral (41.1%) features outweigh connectivity features (15.9%) for the prediction of accelerometry jerk. In contrast, for bar press rate prediction, connectivity (39.8%) emerges as the predominant domain, surpassing spectral (34.7%) and temporal (25.5%) features. Among the individual types of neuro-markers for accelerometry jerk decoding, band power (BP) (33.2%), line length (LL) (21.0%), and band power ratio between channels (BPRC) (10.2%) stand out as the most influential features within the spectral, temporal, and connectivity domains, respectively. This holds true despite the availability of a larger number of connectivity features compared to spectral or temporal features, owing to connectivity’s reliance on the squared number of channels. For bar press rate, BP (18.9%) and BPRC (15.5%) consistently rank as critical, with other spectral and connectivity features like band power ratio between bands (BPRB) (11.2%) and band Pearson correlation (BCorr) (8.2%) also contributing substantially to predictions, unlike other temporal features. Notably, for the leading contributors (BP and BPRC) as well as other neuromarkers that span seven frequency bands, including BCorr, coherence (Coh), and phase locking value (PLV), their high-gamma components are identified as crucial for decoding defensive behaviors, except Coh for accelerometry jerk and PLV for bar press rate, which prominently feature alpha and gamma components, respectively.

**Figure 3.**
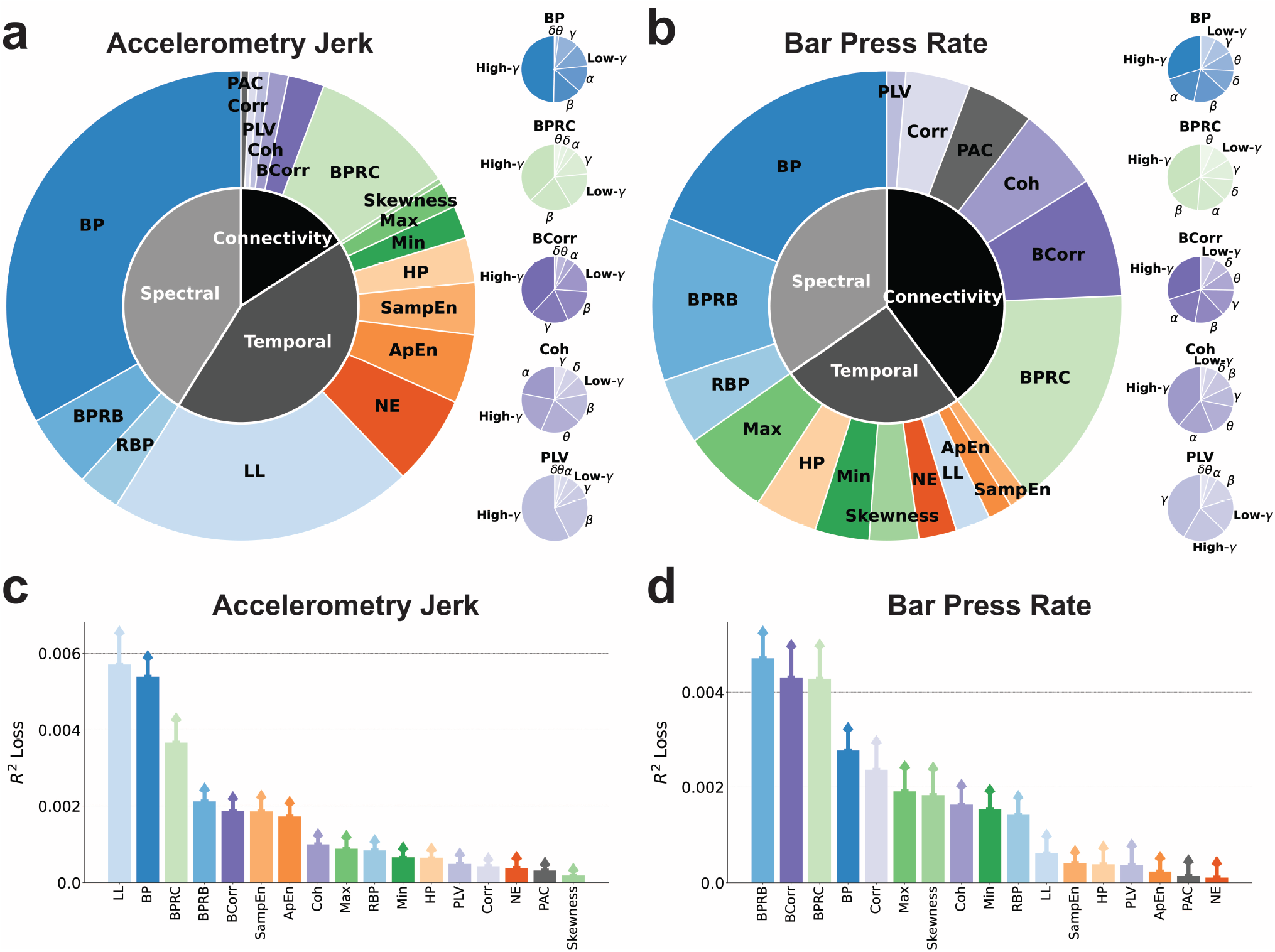
The importance of neuro-markers during decoding process. (a) (Left) The importance of various types of neuro-markers for decoding the accelerometry jerk is illustrated in the outer chart. The importance of markers in the spectral, temporal, and connectivity domains is shown in the inner chart. (Right) The importance of band power, band power ratio between channels, band Pearson correlation, coherence, and the phase locking value in different frequency bands is shown in the small charts. (b) Same as in (a), but for the bar press rate. (c) Contribution of various types of neuro-markers to the decoding performance for accelerometry jerk, averaged across subjects and recording sessions and evaluated using the *R*^2^ loss after removing each type of markers from the ML input. Error bars indicate the standard errors across subjects and sessions. (d) Same as in (c), but for bar press rate.

Beyond quantifying feature importance by evaluating their attribution to the prediction, we also explored their impact on decoding performance, as illustrated in Figures 3(c) and (d). For accelerometry jerk, BP, LL, and BPRC were identified as principal contributors, aligning with their established predictive importance in Figure 3(a). The order of neuro-marker contributions to accelerometry jerk decoding performance as shown in Figure 3(c) mirrors their predictive significance as depicted in Figure 3(a). In the case of bar press rate, BCorr, BPRC, and BPRB maintain a substantial impact on decoding performance, consistent with Figure 3(b). However, BP’s contribution appears noticeably diminished relative to the aforementioned features, underscoring its reduced spectral significance in comparison with BPRB for bar press rate decoding.

This analytical approach to feature importance in both prediction and decoding performance elucidates the substantial importance of neuro-markers across all three domains. BP and BPRC emerge as common key contributors for decoding both defensive behaviors, with LL for accelerometry jerk and BCorr and BPRB for bar press rate also deemed important in terms of prediction and performance.

### 3.3 Importance and contribution of band powers in different frequency bands to the decoding performance

In Figure 3, band power emerged as one of the most influential features. We delved deeper into its importance in terms of prediction and decoding performance across seven frequency bands, including delta, theta, alpha, beta, low-gamma, gamma, and high-gamma, extracted from both IL and BLA, for the decoding of accelerometry jerk and bar press rate, as detailed in Figure 4. The importance matrices in Figures 4(a)-(d) highlight the importance of band power in these frequency bands across recording sessions, brain regions, and targeted behaviors. Collectively, these matrices consistently reveal that high-gamma power in the IL and BLA is more important for predicting behavior than all other frequency bands, and that this is true across different phases of aversive learning and extinction. Figures 4(e)-(h) compare the pairwise similarities among the elements of the importance matrices from Figures 4(a)-(d), examining either the significance of spectral power within identical bands and sessions across different brain regions or in decoding diverse defensive behaviors. These importance matrices exhibit substantial correlation with each other (*r >* 0.61, *p* < 6.0*e −* 4), with the high-gamma components invariably displaying elevated importance values. This pattern suggests that high-gamma power maintains a consistent association with defensive behavior across various contexts.

**Figure 4.**
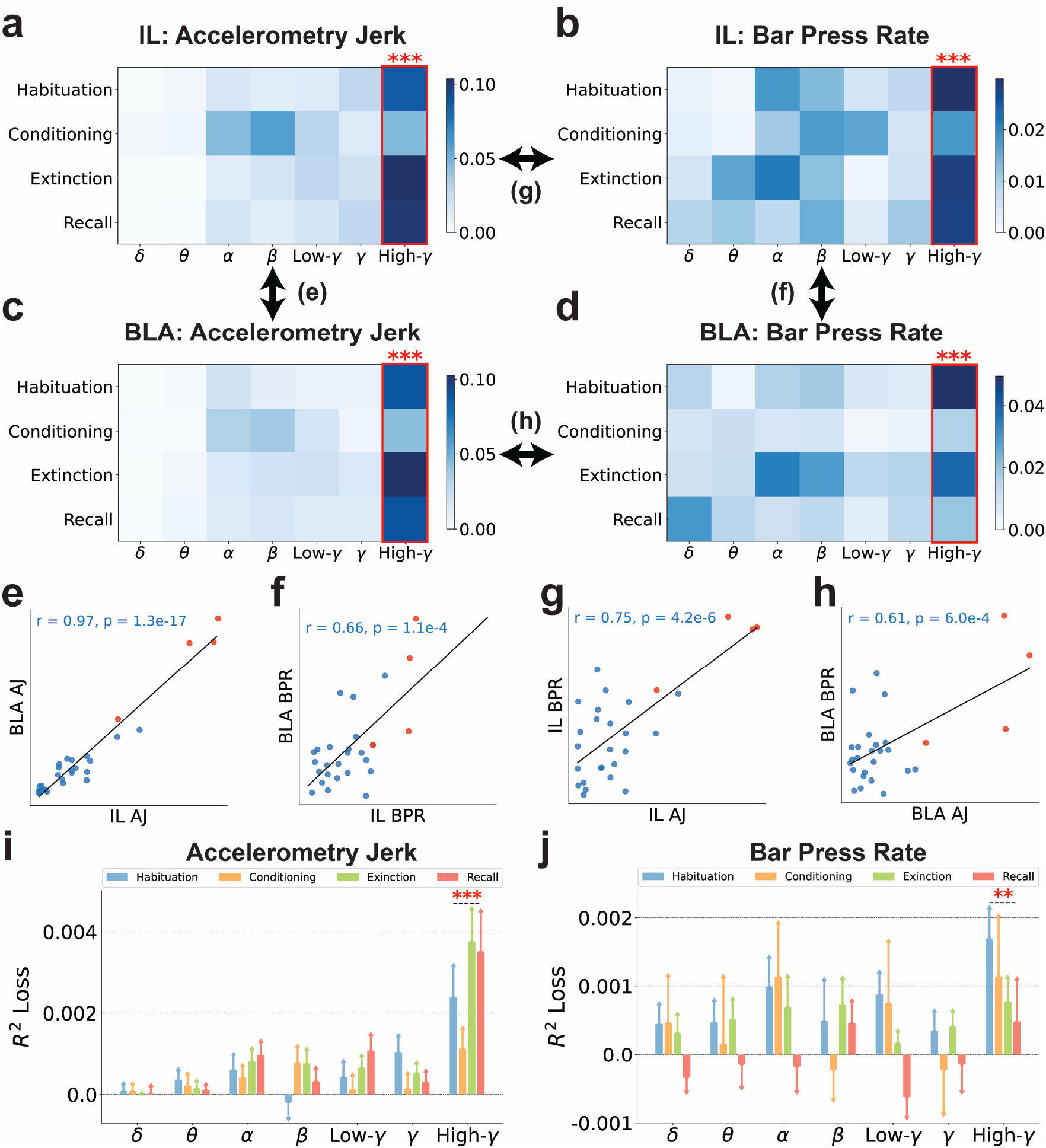
The high-gamma band powers are generally more predictive than other bands for decoding defensive behaviors. (a-d) The importance of band powers (BP) in seven frequency bands and four recording sessions is illustrated in the importance matrices, where each element is the SHAP values averaged across subjects. Bands with significantly higher importance than the other bands are marked with red asterisks. (Independent-sample t-test; *\* * \** : *p* < 0.001). (a) BP in IL for decoding accelerometry jerk. (b) BP in IL for decoding bar press rate. (c) BP in BLA for decoding accelerometry jerk. (d) BP in BLA for decoding bar press rate. (e-h) The similarities between the importance matrices were evaluated using the Pearson correlation coefficient (*r*) and p-value (*p*). Each dot indicates its importance in the same band and the same session between different brain regions or defensive behaviors. Red points denote the importance of high-gamma powers. (e) BP in IL and BLA for decoding accelerometry jerk (AJ). (f) BP in IL and BLA for decoding bar press rate (BPR). (g) BP in IL for decoding AJ and BPR. (h) BP in BLA for decoding AJ and BPR. (i-j) The contribution of BP in seven frequency bands to the decoding performance, averaged across subjects, was evaluated using the *R*^2^ loss after removing each band from the ML inputs. The error bars indicate the standard errors across subjects. Bands with significantly higher contributions than the other bands are marked with asterisks. (Independent-sample t-test; *\*\** : *p* < 0.01, *\*\*\** : *p* < 0.001). (i) The contribution of BP to the decoding performance for accelerometry jerk. (j) The contribution of BP to the decoding performance for bar press rate.

Expanding our analysis to consider the band powers from another angle, we explored their impact on decoding performance across seven frequency bands, as depicted in Figures 4(i) and (j). Notably, the exclusion of high-gamma power leads to a significantly stronger decline in model performance across subjects and sessions compared with all other bands, aligning with observations from Figures 4(a)-(h). Therefore, the comprehensive findings of Figure 4 underscore the pivotal role of high-gamma power as the spectral band most closely linked to defensive behavior, both in terms of attribution to prediction and decoding performance.

### 3.4 Importance and contribution of cross-region neuro-markers in different frequency bands to the decoding performance

In Figure 3, the band power ratio between IL and BLA emerged as a pivotal feature, especially in the context of bar press rate decoding. We dissected the relative contribution of different frequency bands as depicted in Figure 5. Here again, high gamma features were identified as the most influential encoders of defensive behaviors. Additionally, beta band ratios from BLA to IL exhibited marginal significance for accelerometry jerk, as illustrated in Figure 5(c). Figures 5(e)-(h) explore the pairwise similarities among the elements of the importance matrices from Figures 5(a)-(d), assessing either the importance of spectral power ratios within identical bands and sessions across two reciprocal ratios (IL/BLA and BLA/IL) or in decoding various defensive behaviors. These comparisons revealed significant similarities (*r >* 0.67, *p* < 1.1*e −* 4). High-gamma power ratios were distinctly more important than other bands in various analyses presented in Figures 5(e)-(h). This comprehensive analysis indicates a clear concordance in the significance of high-gamma power ratios between IL and BLA, aligning with the patterns of importance outlined in Figures 5(a)-(d).

**Figure 5.**
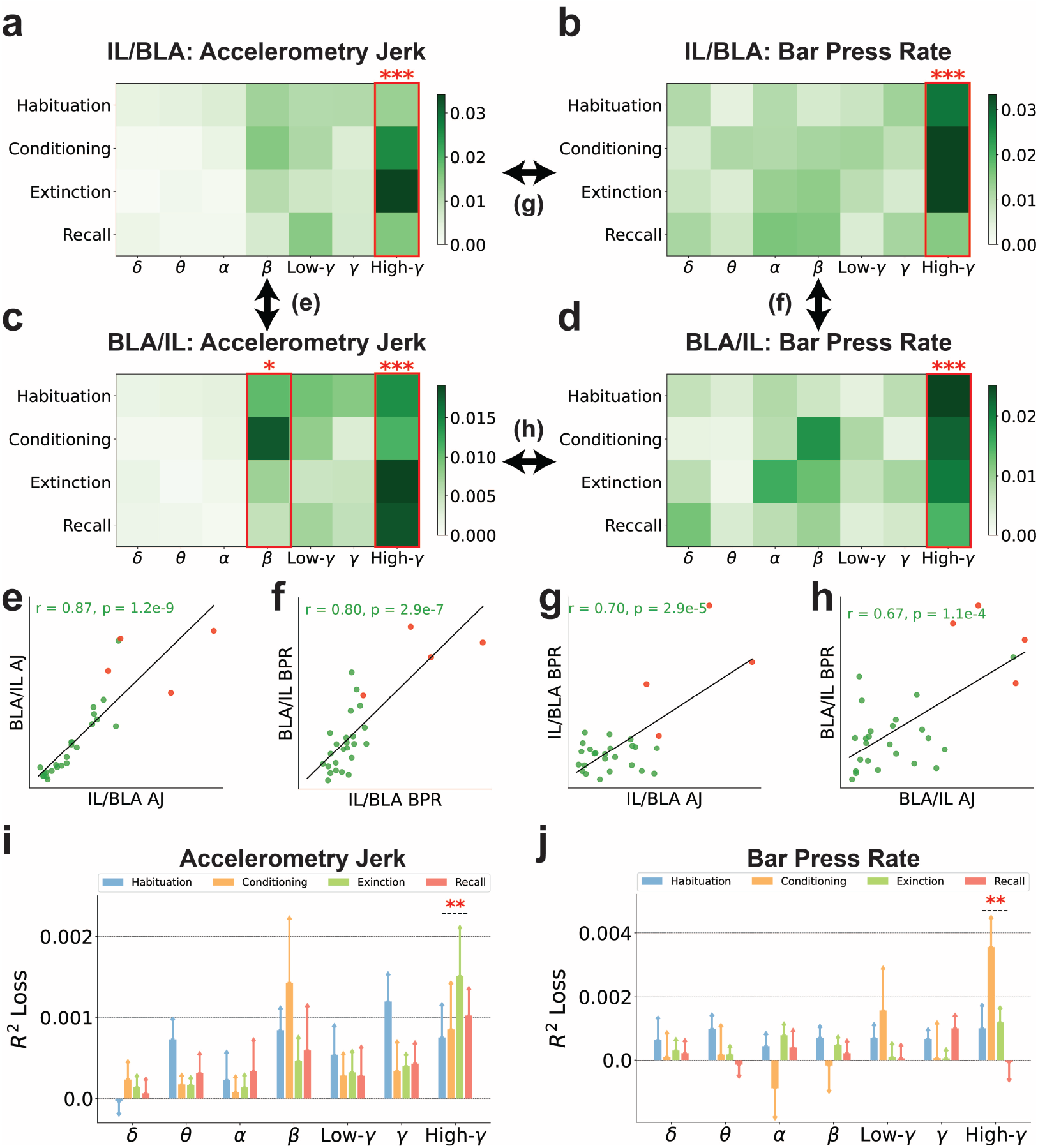
The high-gamma band power ratios between IL and BLA are generally more predictive than other bands for decoding defensive behaviors. (a-d) The importance of band power ratios between IL and BLA (BPRC) in seven frequency bands and four recording sessions is illustrated in the importance matrices, where each element is the SHAP values averaged across subjects. Bands with significantly higher importance than the other bands are marked with red asterisks. (Independent-sample t-test; *\** : *p* < 0.05, *\* * \** : *p* < 0.001). (a) BPRC of IL to BLA (IL/BLA) for decoding accelerometry jerk. (b) IL/BLA for decoding bar press rate. (c) BPRC of BLA to IL (BLA/IL) for decoding accelerometry jerk. (d) BLA/IL for decoding bar press rate. (e-h) The similarities between the importance matrices were evaluated using the Pearson correlation coefficient (*r*) and p-value (*p*). Each dot indicates its importance in the same band and the same session between different brain regions or defensive behaviors. Red points denote the importance of high-gamma power ratios. (e) BPRC of IL to BLA (IL/BLA) and BPRC of BLA to IL (BLA/IL) for decoding accelerometry jerk (AJ). (f) IL/BLA and BLA/IL for decoding bar press rate (BPR). (g) IL/BLA for decoding AJ and BPR. (h) BLA/IL for decoding AJ and BPR. (i-j) The contribution of BPRC in seven frequency bands to the decoding performance, averaged across subjects, was evaluated using the *R*^2^ loss after removing each band from the ML inputs. The error bars indicate the standard errors across subjects. Bands with significantly higher contribution than the other bands are marked with asterisks. (Independent-sample t-test; *\*\** : *p* < 0.01). (i) The contribution of BPRC to the decoding performance for accelerometry jerk. (j) The contribution of BPRC to the decoding performance for bar press rate.

To gain further insights into the band power ratios, we examined their impact on decoding performance, as illustrated in Figures 5(i) and (j). High-gamma power ratios consistently led to the most substantial decrease in performance across subjects and sessions when excluded from the ML model. Thus, high-gamma power ratios are critical to decoding performance for both accelerometry jerk and bar press rate, surpassing the impact of all other frequency bands.

The band power ratios reveal variations in the activation levels between IL and BLA, offering insights into their differential engagement during defensive behaviors. These ratios allow researchers to deduce the degree of synchronization and the dynamic interactions between IL and BLA. However, it is important to note that band power ratios alone do not directly quantify the functional connectivity of these regions. Consequently, we further explored the Pearson correlations between neural signals of IL and BLA, evaluating their significance for prediction and impact on decoding performance across various frequency bands, as depicted in Figure 6. In this analysis, correlations within the high-gamma frequency band emerged as the most informative features, outperforming those of other frequency bands in decoding both accelerometry jerk and bar press rate, as shown in Figures 6(a) and (b). Figure 6(c) demonstrates the similarity between the elements of the importance matrices from Figures 6(a) and (b), revealing a significant correlation (*r* = 0.53, *p* = 3.6*e −* 3). We further explored the impact of band Pearson correlations on decoding performance, as depicted in Figures 6(d) and (e). High-gamma correlations consistently led to the most significant decline in performance when excluded from the model. Collectively, band power ratios and band Pearson correlations elucidate the neural representations between IL and BLA through distinct lenses. Therefore, the findings presented in Figures 4, 5 and 6 together show that, across spectral and connectivity domains, oscillations in the high-gamma range within and between IL and BLA are the most reliable encoder of defensive behaviors. Thus, they may be the most reliable features for closed-loop decoding and intervention.

**Figure 6.**
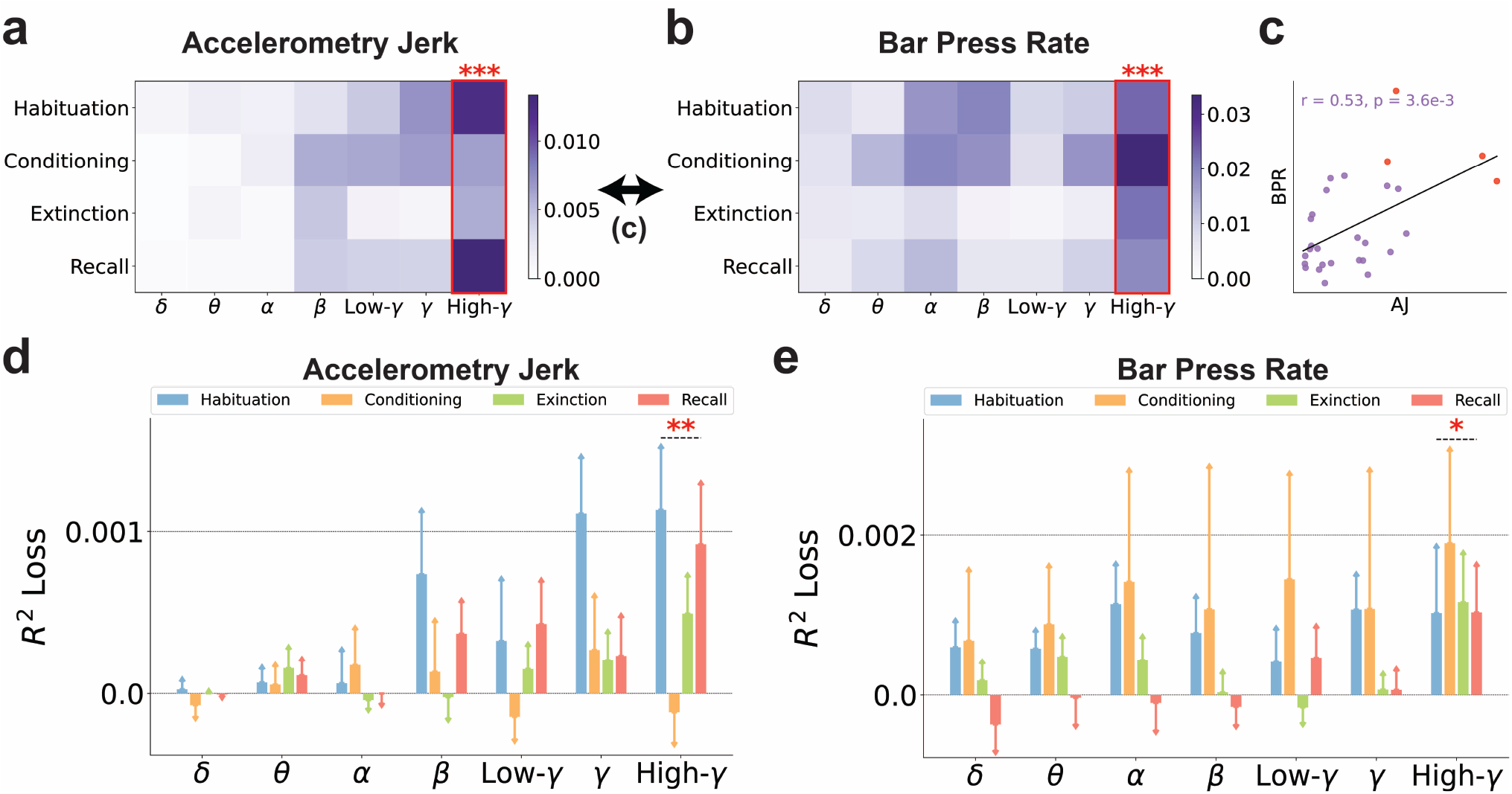
The high-gamma band Pearson correlations between IL and BLA are generally more predictive than other bands for decoding defensive behaviors. (a-b) The importance of band Pearson correlations between IL and BLA (BCorr) in seven frequency bands and four recording sessions is illustrated in the importance matrices, where each element is the SHAP values averaged across subjects. Bands with significantly higher importance than the other bands are marked with red asterisks. (Independent-sample t-test; *\*\*\** : *p* < 0.001). (a) BCorr for decoding accelerometry jerk. (b) BCorr for decoding bar press rate. (c) The similarity between the importance matrices for decoding accelerometry jerk and bar press rate was evaluated using the Pearson correlation coefficient (*r*) and p-value (*p*). Each dot indicates its importance in the same band and the same session between different defensive behaviors. Red points denote the importance of high-gamma correlations. (d-e) The contribution of BCorr in seven frequency bands to the decoding performance, averaged across subjects, was evaluated using the *R*^2^ loss after removing each band in the ML inputs. The error bars indicate the standard errors across subjects. Bands with significantly higher contributions than the other bands are marked with asterisks. (Independentsample t-test; *\** : *p* < 0.05, *\*\** : *p* < 0.01). (d) The contribution of BCorr to the decoding performance for accelerometry jerk. (e) The contribution of BCorr to the decoding performance for bar press rate.

### 3.5 Feature selection chooses important neuro-markers and maintains decoding performance

In this study, we introduced 17 types of neuro-markers as features, yielding a total of 1296 features for inclusion in our ML framework. Incorporating all these features would lead to increased computational and memory demands. Feature selection is a widely recognized strategy for mitigating the computational burden of cognitive decoders [62, 116]. Figure 7 explores the impact of feature dimensionality on decoding performance and the proportion of various types of neuro-markers among the selected features. Figure 7(a) presents the feature selection process based on feature importance as quantified by SHAP values. Here, the decoding accuracy on the validation sets, averaged across subjects, is depicted in relation to the quantity of top-ranked features. Notably, performance improves with an increasing number of selected features, reaching a plateau at approximately 100 features. By selecting only 36 and 81 top-ranked features, we observed that decoding accuracy on validation sets across all recording sessions was comparable to, and not significantly inferior to, the optimal performance identified through an exhaustive exploration of all possible counts of top-ranked features (Paired-sample t-test; accelerometry jerk: *p* = 1.1*e−*1, bar press rate: *p* = 6.1*e −* 2. See section 2.8). These findings underscore the feasibility of dramatically reducing feature dimensionality by 97.2% (36 out of 1296) and 93.8% (81 out of 1296) without significantly compromising decoding efficacy. Within the subset of 36 and 81 top-ranked features selected for the decoding of accelerometry jerk and bar press rate, respectively, an average of 10.7 features are concordant and can predict both defensive behaviors across subjects.

**Figure 7.**
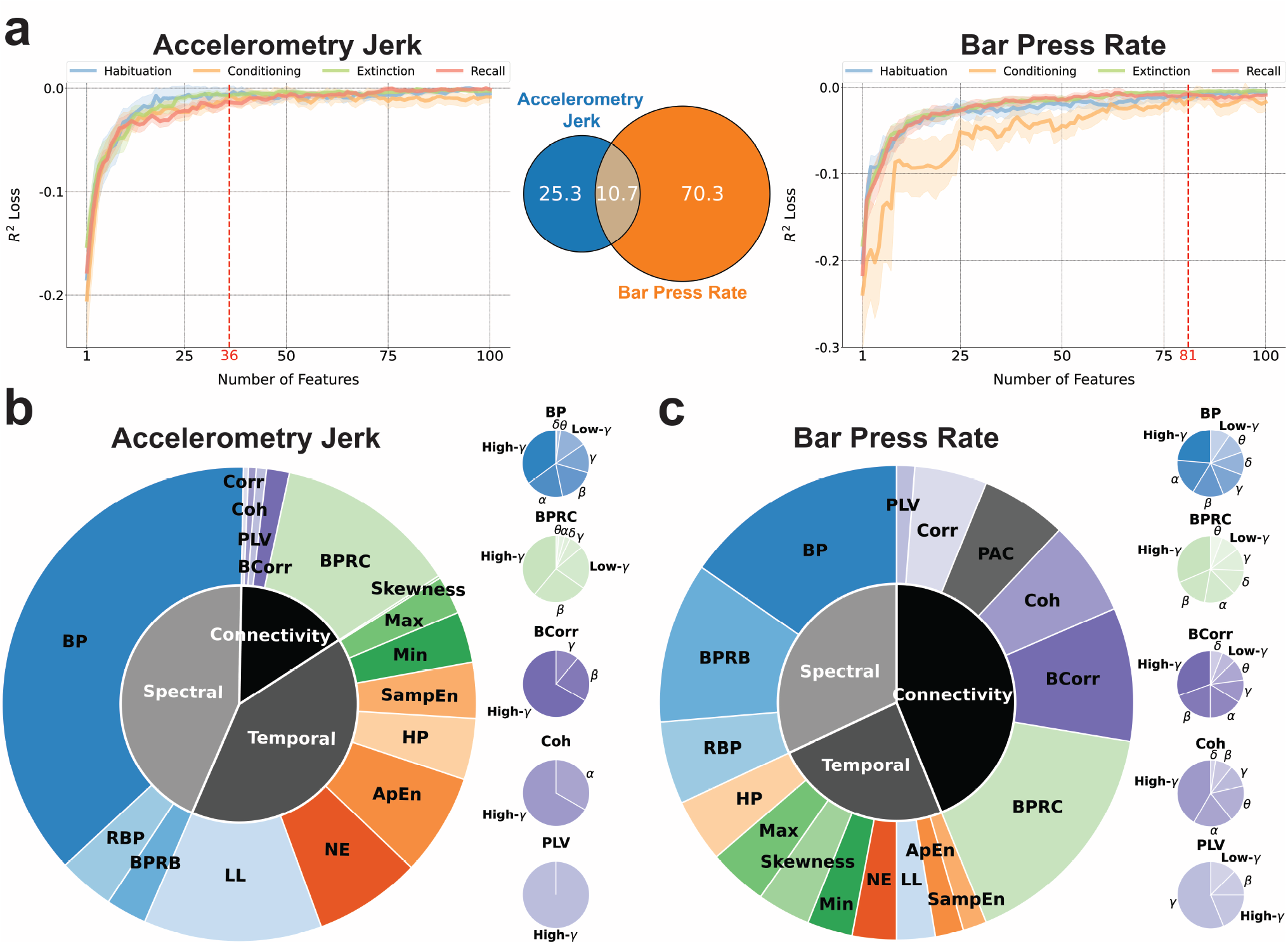
Neuro-markers with higher feature importance and contribution to the decoding performance were chosen through feature selection. (a) Feature selection using feature importance evaluated by SHAP values for decoding the accelerometry jerk (Left) and bar press rate (Right). Features were iteratively selected according to the decreasing order of SHAP feature importance. Shading areas indicate the standard errors across subjects. 36 and 81 top-ranked features were selected from all 1296 features respectively. The performance saturates after 100 features. Venn diagram represents the overlap of top-ranked features for the accelerometry jerk and bar press rate (Middle). (b) (Left) The proportion of different types of neuro-markers among the selected top-ranked features for decoding the accelerometry jerk is illustrated in the outer chart. The proportion of selected features in the spectral, temporal, and connectivity domains is shown in the inner chart. (Right) The proportion of band power, band power ratio between channels, band Pearson correlation, coherence, and the phase locking value in different frequency bands are shown in the small charts. (c) Same as in (b), but for decoding the bar press rate.

Figure 7(b) and (c) delineate the distribution of different types of neuro-markers within the selected features, aligning with the previously established importance of these markers regarding prediction and performance as depicted in Figure 3. In the case of accelerometry jerk, spectral (43.8%) and temporal (40.6%) features were more frequently selected over connectivity features (15.6%). BP (37.2%), LL (12.2%), and BPRC (12.5%) emerged as the predominant feature groups within the spectral, temporal, and connectivity domains, respectively, as shown in Figure 7(b). Conversely, for bar press rate, as illustrated in Figure 7(c), the model exhibited a preference for selecting connectivity features (43.8%) over spectral (31.9%) and temporal (24.2%), with BPRC (16.2%), BP (15.4%), BPRB (11.0%), and BCorr (9.1%) identified as leading predictors. Across all neuro-markers that span 7 frequency bands, including BP, BPRC, BCorr, Coh, and PLV, high-gamma components were most frequently chosen for decoding both defensive behaviors, with the exception of PLV for bar press rate, where the gamma component was more prominently featured.

Since there can be a lag between decisions and manifested behavior, we also evaluated the decoding results using lagged neural data, as presented in Figure A1. Figures A1(a) and (b) illustrate that neural features temporally close to the current time point yield superior decoding accuracy for both behaviors, suggesting these features encapsulate a richer neural representation regarding defensive responses than those from earlier time windows. As shown in Figure A1(c), the inclusion of features from preceding time windows together with current features does not markedly enhance the decoding performance for accelerometry jerk. In contrast, Figure A1(d) indicates a modest improvement in the decoding of bar press rate when previous time window features are incorporated. Furthermore, Figures A1(e) and (f) indicate that the predictive power of features for both behaviors is predominantly concentrated in recent time windows, with a significant decline in predictivity as the temporal gap widens. The findings indicate that neural representations closest to the event of interest are most informative for decoding both defensive behaviors, with immediate past features contributing more significantly to model accuracy than older ones. We thus have emphasized the importance, in preceding and subsequent analyses, of features aligned to behavior with zero lag. The demonstrated temporal gradient in feature predictivity could inform the development of a more refined real-time decoder for neuropsychiatric interventions.

In Figure 8, we explore the dependency of decoding performance on the diverse types of neuro-markers employed and the dimensionality of the feature set. This comparison is made between decoding outcomes utilizing only conventional band powers, decoding with a selected group of features as identified in Figure 7, and decoding with the entire set of extracted features. Figures 8(a)-(d) present the decoding performance for accelerometry jerk and bar press rate using band power features, selected top-ranked features, and all features, assessed by *R*^2^ and *r* metrics, respectively. The addition of other neuro-markers beyond only band power, coupled with feature selection, significantly enhances performance across sessions (for accelerometry jerk, *R*^2^ from 0.4815 to 0.5357, *r* from 0.7229 to 0.7579; for bar press rate, *R*^2^ from 0.3073 to 0.3476, *r* from 0.5708 to 0.6092). Moreover, employing the limited feature set as delineated in Figure 7 does not lead to a significant reduction in performance when compared to the utilization of all features. This observation holds true for the decoding of both defensive behaviors evaluated by both *R*^2^ and *r* metrics. Notably, the adoption of feature selection exceptionally reduces the model’s time complexity, achieving training times of less than 110 ms and inference times of less than 1 ms across all subjects and sessions.

**Figure 8.**
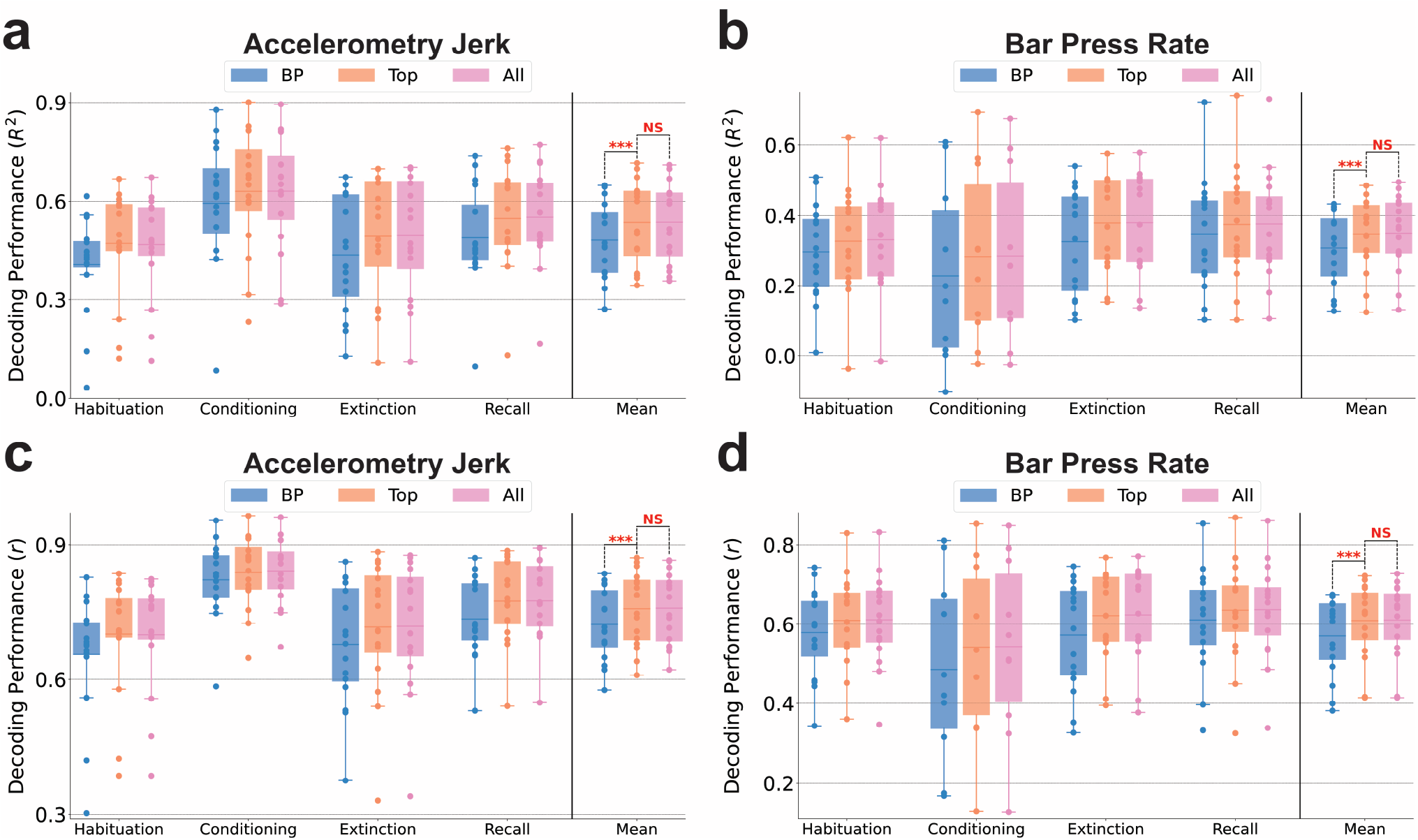
The decoding performance was improved by using more than one type of neuro-marker, but remained good when only high-importance features were used for training. (a) The comparisons of decoding performance for the accelerometry jerk using band powers, selected top-ranked features, and all features in four recording sessions averaged across subjects, evaluated using the coefficient of determination (*R*^2^). Mean denotes the performance averaged across sessions. The line in the boxplot shows the average performance across subjects, and each dot indicates the result of an individual subject. The comparisons in Mean were marked with asterisks if one averaged performance was significantly greater than the other. (Wilcoxon signed-rank test; *\*\*\**: *p* < 0.001, *NS* : *p ≥* 0.05) (b) Same as in (a), but for decoding the bar press rate. (c) Same as in (a), but evaluated using the Pearson correlation coefficient (*r*). (d) Same as in (c), but for decoding the bar press rate.

Collectively, the results presented in Figures 7 and 8 underscore that the feature selection process effectively identifies important features with significant additive attribution to the prediction output and remarkable contribution to the performance in decoding defensive behaviors. Through this process, a select group of top-ranked features not only sustains decoding performance with remarkably reduced time complexity but also enhances performance in comparison to relying exclusively on band power features.

## 4 Discussion

We developed a machine learning framework for accurately decoding defensive behaviors from multi-channel local field potentials recorded from the infralimbic cortex and basolateral amygdala. Critically, accelerometry jerk and bar press rate exhibited higher decodability compared to the freezing score, as evidenced by both the training dynamics and performance evaluations on the test set (Figure 2). These two decodable behaviors were encoded by distinct sets of highly informative features (Figures 3 and 7).

This research builds upon our previous work, which underscored that these metrics each capture unique facets of defensive behavior [83]. The variation in encoding between behaviors suggests that they may have distinct neural substrates, i.e., that a closed-loop system designed to modulate defensive processes might need to control different aspects of cortical/amygdala physiology depending on the exact process being targeted. The challenge in accurately decoding the freezing score — conceptually the inverse of freezing and calculated from video frame changes to approximate the rat’s horizontal velocity — is intriguing. Given its mathematical relationship with accelerometry jerk, which essentially represents a higher derivative of movement than freezing score, this difficulty is unexpected. On the other hand, considering that mammalian motor control often optimizes for minimum jerk [117], it stands to reason that such dynamics are more directly encoded in the neural circuitry that eventually affects motor planning.

Freezing, as derived from the freezing score, is probably the single most common behavior used to study the IL, BLA, and the broader circuits of the extended amygdala [79, 118, 119]. Its association with various LFP processes, particularly emphasizing local oscillations and cross-regional synchrony within the theta band, is well-documented [75,86]. Therefore, our inability to decode this behavior accurately presents a notable discrepancy. One possible explanation for this difference could be our focus on decoding second-to-second changes in behavior, in contrast to previous studies that typically examined longer timescales, such as the percentage of a cue tone spent in freezing versus other behaviors. As illustrated in Figure 2(c), our decoders demonstrated better performance in capturing these broader timescales (trends or global means) than short-term variability in freezing score. This observation aligns with our previous behavioral research, which indicated that the mean freezing score across subjects correlated more closely with the mean accelerometry jerk and bar press rate, than when analyzing individual subjects [83]. This may be attributed to the averaging process across subjects, which effectively smoothed away local variance while preserving global trends, thereby rendering freezing score more comparable with other measured behaviors.

Beyond the conventional use of band power features for decoding cognitive and emotional processes [38, 62, 66], our model incorporates a broader array of neuro-markers across spectral, temporal, and connectivity domains. Temporal features demonstrated a particularly significant contribution to decoding accelerometry jerk over bar press rate, as evidenced in Figures 3(a) and (b). This disparity likely stems from the capability of temporal-domain features to capture changes over very short intervals, reflecting the dynamic and swift variations in the defensive behaviors that define the accelerometry jerk data. Interestingly, connectivity features played a more pronounced role in decoding bar press rate compared to accelerometry jerk. This distinction may reflect the difference in behavioral characterization underpinning these behaviors; unlike accelerometry jerk, bar press rate involves the suppression of a reward-seeking response, diverging from motion-based defensive behaviors like freezing. Hence, prior studies linking defensive behaviors with theta oscillations and cross-regional LFP connectivity may more accurately depict variations in reward-related processes. A noteworthy finding is that, alongside coherence (Coh), significant decoding insights were derived from the band power ratio between channels (BPRC) and band Pearson correlation (BCorr). Thus, BPRC and BCorr warrant increased consideration over Coh in subsequent fear regulation research. We have demonstrated that these features encompass unique information not captured by band power alone [38, 67, 103, 120].

The high-gamma band was particularly important for decoding accelerometry jerk and bar press rate in BP, BPRC, and BCorr (Figures 4-6). This finding contrasts with earlier research, where fear-related behavior was primarily correlated with theta band power and sycnhrony [75, 86]. The divergence in findings could stem from our distinct analytical methodology. Whereas previous studies often explored categorical differences, such as contrasting animals showing low versus high freezing behavior in a dichotomized analysis, our approach aimed to directly predict behaviors within individual animals and sessions. Within this shorter timescale, the involvement of faster processes, like those within the high-gamma range, may become more pivotal. We also used different electrodes, with tighter spacing that emphasizes local signals within IL and BLA. This again would emphasize more spatially local high-frequency components over more spatially distributed low-frequency LFPs. However, this emphasis on local signals more realistically models a clinical scenario, where electrodes would be implanted within a relatively small brain region.

Through our feature selection process, we strategically chose a limited subset of features to minimize the computational complexity and memory demands of our ML framework. Utilizing only 36 and 81 top-ranked features, as depicted in Figure 7, we not only significantly surpassed the decoding performance achieved with 56 BP features but also matched the performance obtained with the full set of 1296 features, as demonstrated in Figure 8. This indicates that neuro-markers other than BP encode unique information critical for decoding. The analytical findings from Figures 3-6 further support that incorporating a broader spectrum of neural representations enhances decoding effectiveness, offering a more nuanced insight into neuro-markers’ roles in modulating defensive behaviors. Additionally, our results imply the existence of a considerable number of features that are either non-predictive or redundant within the model. The feature selection process effectively eliminates less informative features for each subject, thereby significantly reducing computational expenses during training and inference phases and lowering memory requirements. These efficiencies, combined with the high decoding accuracy, underscore the importance of an optimized feature selection strategy for neural decoders in neuropsychiatric brain-machine interfaces (BMIs).

Advanced machine learning models have been shown to markedly enhance neural decoding performance over conventional approaches. In our investigation, we assessed the decoding capabilities of state-of-the-art models in neural decoding tasks, including linear regression (LR), support vector machine with linear (SVM-Lin) and radial basis function kernels (SVM-RBF), random forest (RF), multilayer perceptron (MLP), long short-term memory (LSTM), convolutional neural networks (CNN), and Light Gradient Boosting Machine (LightGBM). Building on our prior research on seizure detection [90, 91, 121], mental fatigue prediction [68], finger movement classification [107,122], and tremor detection from electrophysiological signals [97,123], gradient-boosted decision tree models (GBDT) including LightGBM were found to outperform traditional ML models, including SVM and linear discriminant analysis (LDA). Our findings further reveal that LightGBM was the best-performing model in 14 out of 16 comparisons across decoding tasks, for accelerometry jerk and bar press rate across four recording sessions, as evaluated by the coefficient of determination (*R*^2^) and the Pearson correlation coefficient (*r*). Although RF performed slightly better than LightGBM in decoding accelerometry jerk during the conditioning session as per *R*^2^ and in decoding bar press rate as per *r*, LightGBM demonstrated significantly shorter training (accelerometry jerk: <508 ms, bar press rate: <487 ms) and inference times (accelerometry jerk: <0.8 ms, bar press rate: <0.8 ms) compared to RF (training times: accelerometry jerk: <40 s, bar press rate: <130 s; inference times: accelerometry jerk: <19 ms, bar press rate: <30 ms). These results underscore LightGBM’s capability to deliver both precise decoding outcomes and remarkably rapid decoding speeds with the utilized neuro-markers, which are the key qualities for a decoder within a closed-loop BMI system. Furthermore, LightGBM offers additional advantages over other ML models: it supports parallel computation, greatly speeding up training and inference processes. Importantly, it exhibits low hardware complexity, as demonstrated in recent low-power hardware implementations of closedloop neuromodulation systems [90, 98, 124]. Collectively, these attributes underscore LightGBM’s potential applicability in future fully-implantabe and closedloop psychiatric BMIs.

A notable limitation of our study is the confinement of neural signals to pre-defined neuro-markers. While these neuro-markers intuitively describe neural representations in an interpretable manner, this approach may overlook critical information present in raw neural signals. In future research, we plan to harness the capabilities of artificial neural networks (ANN) for nonlinear and complex modeling, potentially uncovering hidden neural representations not captured by conventional neuro-markers. Recurrent neural networks such as LSTM could be employed to identify concealed temporal dependencies, while CNN or self-attention mechanisms might be utilized to decode the mixed spatial and temporal information. Despite the superior performance of LightGBM over MLP, LSTM, and CNN in our current analyses, pursuing further investigations into ensemble methods that integrate LightGBM with ANNs, as well as adopting ANN-derived neural representations for LightGBM decoding, represent promising avenues. These approaches could significantly enhance decoding accuracy and the generalization capacity of our models.

In our study, we employed two metrics to assess feature importance: SHapley Additive exPlanations (SHAP) and the loss of *R*^2^ upon feature removal. While SHAP values elucidate each feature’s additive attribution to prediction, they do not explicitly evaluate the necessity of features for decoding performance. Conversely, the loss of *R*^2^ quantifies a feature’s impact on performance, yet this metric might yield ambiguous interpretations in cases of high feature correlation. Additionally, it fails to satisfy the three desirable properties of additive feature attribution methods outlined by SHAP, namely local accuracy, missingness, and consistency [115]. Thus, there is a compelling opportunity for researchers to explore alternative metrics for evaluating feature importance in terms of prediction and performance that both minimize computational complexity and embody the aforementioned properties. These metrics also highlight a specific limitation of the LightGBM approach: although we can identify which bands/features are most important for a given analysis (here, high-gamma), we cannot directly use that importance for a simple, biomarker-driven intervention. Tree-based methods focus on dichotomizing a given feature at a specific value, but can select that feature again at deeper tree levels if needed. Thus, they can model complex nonlinear and non-smooth relationships between neural signals and behavior. Unlike a simpler model such as a linear regression, however, tree-based methods do not produce clear or simple relations such as “to decrease defensive behaviors, it would be desirable to reduce BLA high-gamma power”. Inferring and testing such potential causalities would require different approaches, e.g., permuting the model’s inputs in a systematic way and measuring the outputs. On the other hand, the superior decoding accuracy, feasibility for hardware implementation, and substantial pruning potential of tree-based models, as demonstrated in [91, 98, 121] could enable more efficient and effective closed-loop interventions compared to conventional approaches that rely solely on individual biomarkers [125, 126].

In this research, we evaluated our model using an offline paradigm on a dataset aimed at decoding defensive behaviors. To ascertain the robustness of our model across a wider array of neuropsychiatric applications, it would be beneficial to validate our model design using additional datasets, encompassing either identical or divergent tasks. Moreover, transitioning from offline to online neural decoding represents a significant challenge. In our future work, we intend to deploy our decoding framework within an online paradigm, thereby facilitating an assessment of our model’s performance in real-time applications.

## 5 Conclusion

In this study, we analyzed LFP signals from IL and BLA of rats subjected to a tone-shock protocol to extract neuro-markers. These markers were subsequently utilized in our ML decoding framework, which incorporates SHAP-based feature selection and LightGBM for decoding defensive behaviors. Notably, the accelerometry jerk and bar press rate proved to be more decodable than the freezing score. We achieved an average decoding performance of *R*^2^ = 0.5357 and *r* = 0.7579 for the accelerometry jerk, and *R*^2^ = 0.3476 and *r* = 0.6092 for the bar press rate, with exceptionally low time complexity: less than 110 ms for training and less than 1 ms for inference. BP and BPRC emerged as significant neuromarkers for prediction and decoding performance. The high-gamma band within BP, BPRC, and BCorr was consistently identified as crucial for decoding both defensive behaviors across both brain regions. The selection of top-ranked features not only surpassed the performance achieved using only BP features but also maintained the performance level of models utilizing the entire feature set. Our findings underscore the efficacy of developing an accurate and low-latency model for decoding defensive behavior based on LFP features from circuits strongly linked to these behaviors. This work lays the groundwork for future development of an implantable closed-loop psychiatric BMI, showcasing the potential of our framework in advancing neuropsychiatric treatment modalities.

## Acknowledgment

This work was supported by the National Institute of Mental Health Grant R01-MH-123634.

## Ethical statement

All experimental details were approved by the Institutional Animal Care and Use Committee at the University of Minnesota and performed in compliance with the Guide for the Care and Use of Animals. Research facilities were accredited by the American Association for the Accreditation of Laboratory Animal Care.

## Appendix

**Figure A1:**
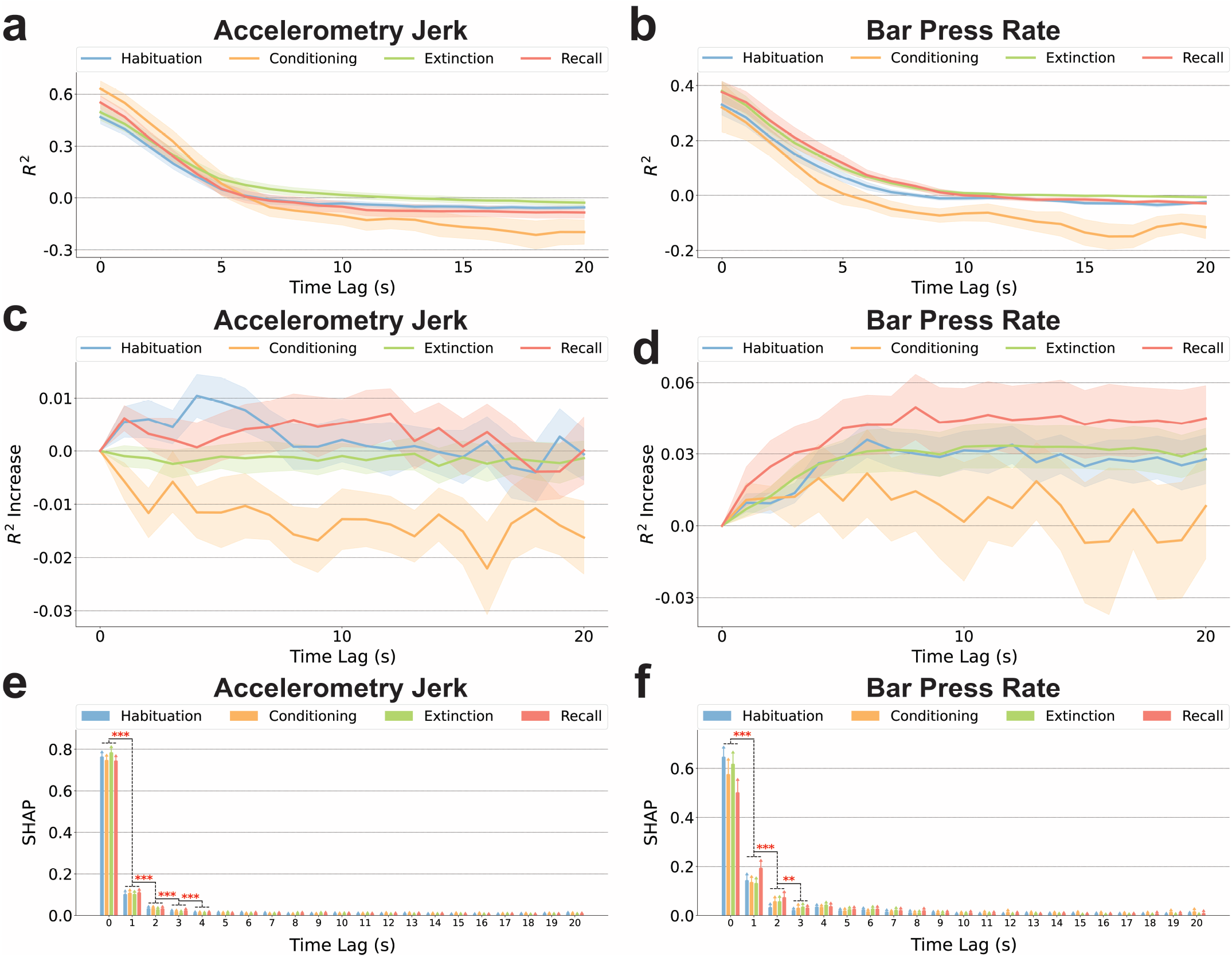
Time-dependent decoding performance and feature importance for two defensive behaviors. (a) Decoding performance for accelerometry jerk using neural features either aligned at zero lag to behavior or lagged by up to 20 seconds with a step size of 1 second for each time lag, in four recording sessions averaged across subjects, evaluated using *R*^2^. The shaded areas indicate the standard errors across subjects. (b) Same as (a), but for decoding bar press rate. (c) Decoding performance for accelerometry jerk when using features from more than one time lags, assessed using an increase in the *R*^2^ compared to a baseline without temporal integration, in four recording sessions averaged across subjects. Neural features from both the current and preceding time windows, extending up to 20 seconds, were incorporated. The shaded areas indicate the standard errors across subjects. (d) Same as (c), but for decoding bar press rate. (e) SHAP importance of features integrating data from the current and preceding time windows up to 20 seconds, averaged across subjects in four sessions for decoding accelerometry jerk. The error bars indicate the standard errors across subjects. The comparisons between features from consecutive time windows are marked with asterisks if one time window’s SHAP values are significantly greater than those of the preceding window across sessions and subjects (Paired-sample t-test; *\*\** : *p <* 0.01, *\* * \** : *p <* 0.001). (f) Same as (e), but for decoding bar press rate.

